# Stable, Variable, Encoding: Distinct Roles of SST, VIP, and EXC Neurons in Visual Novelty Processing

**DOI:** 10.64898/2026.04.09.717494

**Authors:** Karni Lev Bar-Or, Vyshnavi Sankaran Krishnan, David Walker Gauthier

## Abstract

Detecting and processing novelty is critical for learning and survival, yet the stability and flexibility of novelty representations at the level of single neurons remain poorly understood. How novelty evoked responses persist across time, whether novel stimuli are encoded in a stimulus-specific or non-specific manner, and how encoding adapts under changing conditions remain largely unknown. Importantly, novelty responses involve both excitatory and inhibitory neurons, highlighting the need to understand how these cell types differentially contribute to stable and flexible cortical representations. We analyzed longitudinal calcium imaging dataset from mouse visual cortex, tracking excitatory (EXC), somatostatin-expressing (SST), and vasoactive intestinal peptide-expressing (VIP) neurons across six days of a change detection task incorporating contextual novelty, stimulus omissions, and absolute novelty. At the population level, novelty responses were stable across days. However, single-neuron analysis revealed marked instability in EXC and VIP neurons. SST neurons exhibited the highest single-cell stability across all conditions, suggesting a role in maintaining consistent sensory representations. VIP neurons displayed stable responses only to omissions. Regarding information content of novelty responses, we found that EXC neurons encoded both stimulus-specific and non-specific novelty while VIP neurons uniquely transitioned from non-specific to mixed encoding under absolute novelty, revealing previously unrecognized flexibility. These findings reveal distinct, cell-type-specific roles in novelty processing, with SST cells supporting stability and VIP cells adapting their coding to novelty type.

## Introduction

Detecting novelty is crucial for survival and underlies learning and behavioral adaptation in dynamic environments [1,2]. Novelty is defined as a deviation from learned expectations, and its detection depends on prior learning of familiar stimuli and on stimulus identification. Across sensory modalities, novel or unexpected stimuli elicit enhanced neural responses relative to expected ones [1–4], often triggering changes in attention, behavior, and the initiation of learning and memory formation [1,2,5,6]. Importantly, novelty responses are not uniform. They vary by timescale and the form of deviation. Three broad novelty types have been described: contextual novelty, where a previously observed stimulus is novel in the context of recently observed stimuli and thus deviates from the local stimulus context [7–9]; stimulus omission, where an anticipated stimulus is absent from an expected predictable sequence [10]; and absolute novelty, involving exposure to a stimulus that has not been experienced before [11].

In the sensory cortex, both excitatory and inhibitory neuronal populations contribute distinctively to novelty processing [12–14]. In primary visual cortex (V1), excitatory pyramidal neurons (EXC) exhibit increased activity in response to novel stimuli [15–23]. Vasoactive intestinal peptide (VIP) expressing interneurons respond to both novel stimuli and omissions [15,22], while somatostatin-expressing (SST) interneurons are required for contextual novelty detection [18], and show increased activity to familiar stimuli [24]. These findings support disinhibitory circuit models, wherein VIP neurons enhance EXC activity by suppressing SST cells in response to novel stimuli [25–29]. Yet, the distinct roles of these cell types across different novelty conditions—and their contributions over time—remain unclear.

A central open question is whether novelty representations are stable across time, and whether distinct cell types contribute differentially to the stability and flexibility of these responses at the single-cell level. Although cortical circuits are inherently dynamic, consistent sensory perception requires that neuronal circuits maintain stable representations of sensory input over time [30–34]. Previous work has demonstrated robust novelty responses, but these studies were largely restricted to single-day measurements [2,35–37], leaving unresolved their long-term consistency. Key questions remain: First, are population-level novelty responses stable across days and cell types during high function task of novelty detection? Second, if population-level stability is observed, does it reflect consistent responses in individual neurons, or stable population-level dynamics despite individual variability? Evidence on the stability of neuronal representations is mixed—some studies report that individual neurons maintain stable activity over time [38,39], while others argue that stability emerges only at the population level, despite single-cell variability. The latter view is grounded in longitudinal recordings from hippocampus, piriform, barrel, parietal, auditory, and visual cortices, which show that stable population codes can arise from circuits composed of individually unstable neurons [40–51]. However, these studies have primarily focused on familiar stimuli and excitatory cells. Whether such volatility also characterizes novelty responses remains unknown. Third, do cell types differ in single-neuron stability? Elucidating this is essential for understanding how excitatory and inhibitory cells interact to generate robust novelty responses.

Concurrently, the nature of the information content encoded by novelty responses remains under debate. Do novelty responsive neurons signal only the presence of deviance—a non-specific surprise—or do they also encode the identity of the novel stimulus [52–56]? Some studies suggest that only highly stimulus selective excitatory neurons amplify responses to unexpected stimuli [22], whereas others report enhancement across both selective and non-selective cells [23]. For VIP and SST inhibitory neurons, whether novelty responses are stimulus-specific or non-specific remains uncharacterized. Furthermore, it is unknown whether the type of information encoded by each cell type remains fixed under broader novelty conditions, such as absolute novelty. Understanding whether encoding is static or dynamic is crucial, as a change in stimulus encoding might indicate initiation of learning processes.

To address these questions, we analyzed a publicly available longitudinal dataset from the Allen Brain Institute, which tracked EXC, SST, and VIP neurons in mouse visual cortex across six days incorporating a change detection task (contextual novelty), stimulus omissions, and absolute novelty. Notably, this dataset has recently served as the basis for several studies that yielded key insights into differences in the responses of EXC, VIP, and SST cells across novelty conditions, as well as into the types of information encoded by distinct neuronal populations [15,19,20]. Importantly, however, these analyses were restricted to discrete days rather than across all consecutive days of the experiment, and therefore did not address the stability of novelty responses at the level of single neurons, nor coding flexibility. This unique dataset thus enabled us to systematically investigate population-level and single-cell response stability and stimulus specificity across multiple novelty types in genetically defined cell types.

We found that, at the population level, novelty responses were stable across days, with consistent proportions of responsive neurons and similar response magnitudes. However, analysis at the single-neuron level revealed substantial fraction of unstable neurons, indicating that population-level stability is comprised of neurons with unstable responsiveness. SST interneurons exhibited the highest degree of single-cell response stability, suggesting a distinct role in maintaining consistent representations. Notably, VIP neurons displayed stable responses only to omissions. These findings suggest that population-level stability can arise from dynamic, cell-type-specific contributions and that SST neurons may play key roles in stabilizing cortical representations of novelty.

With regard to information content, cell types exhibited distinct contextual novelty responses. EXC neurons displayed both stimulus-specific and non-specific novelty responses. SST neurons exhibited exclusively stimulus-specific responses. VIP neurons, in contrast, showed non-specific responses under familiar conditions, but displayed both specific and non-specific encoding under absolute novelty. This shift suggests that VIP neurons can dynamically modify their coding strategy in response to absolute novelty, pointing to a previously unrecognized flexibility in their function. Supporting this, analysis of a complementary Neuropixels dataset revealed a shift in VIP firing patterns and stimulus-specific responses under absolute novelty. Together, our findings demonstrate that the stability and information content of novelty responses are cell-type specific, highlighting distinct and flexible roles of excitatory and inhibitory populations in cortical novelty encoding.

## Results

### Change detection task

We analyzed datasets from the Allen Brain Institute’s Visual Behavior experiments, which involved calcium imaging of neuronal responses in the mouse visual cortex during a visual change-detection task. The data included longitudinal recordings from mice expressing the calcium indicator GCaMP6 in excitatory neurons, somatostatin-expressing inhibitory neurons, or vasoactive intestinal peptide-expressing inhibitory neurons.

In these experiments, mice were trained on a visual task requiring detection of unexpected image changes (contextual novelty) within sequences of repeatedly presented images drawn from a set of eight stimuli. Upon detecting such a change, mice reported their perception by licking a reward spout. Two-photon calcium imaging was performed across six consecutive experimental days. During the first three days, mice performed the task with familiar images that had been used in the preceding training. Additionally, in 5% of trials without image changes, stimuli were randomly omitted, introducing stimulus omission as another form of novelty. On Day 4, the image set was replaced with entirely novel images, unfamiliar to the mice, thereby introducing a condition of absolute novelty. This novel set remained in use on Days 5 and 6. The experimental design included sessions without reward delivery (passive sessions) on Days 2 and 5, while the remaining sessions were reward-delivering sessions (active sessions) (Figures 1A-1B). Behavioral metrics, including task performance, lick rate, running speed, and pupil diameter, were previously reported by the Allen Institute research group to remain stable across all experimental days [19].

**Fig 1.**
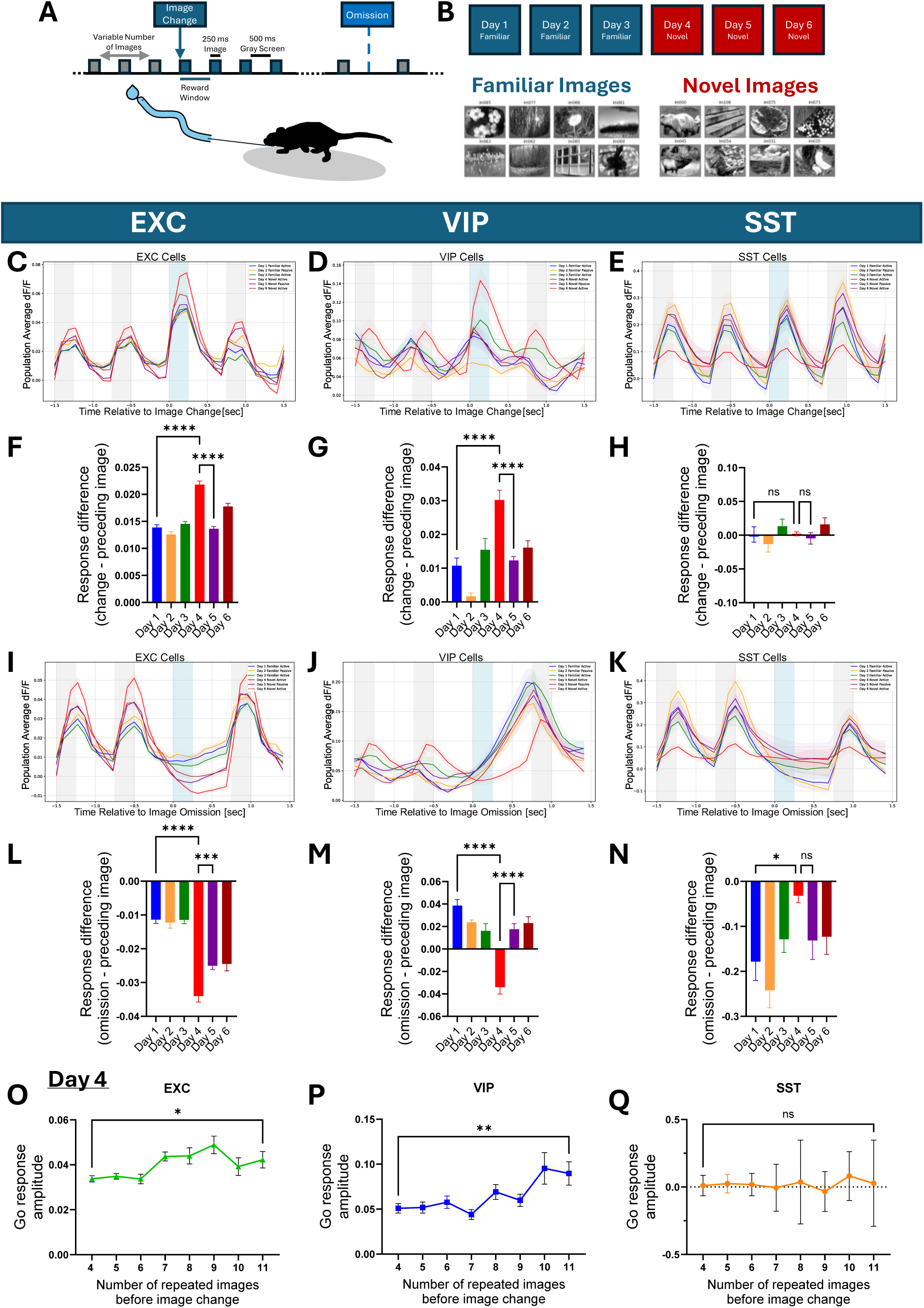
Cell type-specific responses to different novelty conditions. (A) Mice performed a visual change detection task, where natural scene images were repeatedly presented until an image change occurred. In 5% of non-change trials, the stimulus was omitted. (B) In vivo two-photon calcium imaging was performed in the visual cortex across six consecutive days, during which mice performed the task with familiar images on Days 1-3 and with a novel image set on Days 4-6. Days 2 and 5 were passive (non-rewarded) sessions, while the other days were active. (C-E) Population-average responses (ΔF/F) to image presentations (gray) and image changes (Go, blue) for EXC (n = 3124 ± 86 neurons/day), VIP (n = 400 ± 8), and SST (n = 104 ± 1) neurons across all days. (F-H) Go response magnitude (difference between image change and preceding image). EXC and VIP cells showed increased responses on Day 4 (one-way ANOVA followed by Dunnett’s post hoc comparison to Day 4; EXC: F(5, 18739) = 42.2, p < 0.0001; all multiple comparisons p < 0.0001; VIP: F(5, 2393) = 15.5, p < 0.0001; multiple comparisons, p < 0.0001, SST: F(5, 618) = 1.3, p = 0.3; multiple comparisons were ns). (I-K) Population-average responses to stimulus omissions (blue) across days. (L-N) Omission response magnitude (difference between omission and preceding image). EXC and SST neurons showed suppression during omissions, whereas VIP showed increase (one-way ANOVA followed by Dunnett’s post hoc; EXC: F(5, 18,739) = 39.3, p < 0.0001; multiple comparisons were significant, p < 0.0001; VIP: F(5, 2393) = 20.7, p < 0.0001; multiple comparisons were significant, p < 0.0001; SST: F(5, 618) = 3.8, p = 0.002; significant difference for Day4 Vs. Day 1, p = 0.02). (O-Q) Go response amplitude on Day 4 as a function of the number of repeated images prior to image change. EXC and VIP neurons showed increased responses with more image repetitions, consistent with increased contextual novelty (comparison of responses following 4 vs. 11 image repetitions; unpaired t-test; EXC: t(6010) = 2.1, p = 0.03; VIP: t(726) = 2.7, p = 0.006; SST: t(198) = 0.5, p = 0.6). Error bars: ±SEM across neurons. ns p>0.05, *p < 0.05, **p < 0.01, ***p< 0.001, ****p<0.0001.

Using this experimental dataset, we analyzed neuronal activity (ΔF/F fluorescence signals) across multiple days and conditions, quantifying responses from 3124 ± 86 excitatory neurons (N = 7 mice), 104 ± 1 SST neurons (N = 3 mice), and 400 ± 8 VIP neurons (N = 5 mice) per day.

### Cell type-specific responses to different novelty conditions

We first aimed to determine whether stimulus novelty modulates sensory responses across cell types. To this end, we computed average stimulus-triggered population responses across days and conditions, quantifying responses as changes in calcium fluorescence. Image change (hereafter referred to as “Go” responses) and omission response magnitudes were defined as the difference between the response to the image change (or omission) and the response to the immediately preceding image (Figures 1C-1N). Excitatory cells exhibited a characteristic response to image presentations, with a heightened response to image changes [15,19]. The Go response magnitude was significantly increased on Day 4, when a novel image set was introduced, and returned to the baseline levels by Day 5, indicating a transient enhancement by absolute novelty (Figures 1C, 1F). SST cells displayed robust responses to image presentations but showed no increase during the Go. On Day 4, their activity decreased across both image and Go conditions (Figures 1E, 1H), suggesting reduced activity under absolute novelty exposure. VIP cells exhibited a distinct response pattern. On all days except Day 4, VIP activity gradually increased during the interstimulus grey period, peaking before image onset. On Day 4, however, VIP responses were delayed, with peak activity occurring after image onset, and the Go responses were significantly elevated (Figures 1D, 1G). By Day 5 the differences renormalized. Notably, both EXC and VIP cells showed similar modulation on Day 4.

Cell-type differences were also evident during stimulus omissions. When averaged across the population, EXC and SST cells were suppressed during image omissions, whereas VIP cells displayed ramping activity that peaked just before the next image (Figures 1I-1N). VIP omission responses were reduced on Day 4. These results indicate that novelty modulates sensory responses in a cell type-specific manner, consistent with prior studies [15,19,20].

### Dependence of novelty responses on the number of image repetitions preceding a change

The number of images preceding an image change was drawn from a geometric distribution, with most trials containing four repetitions and fewer trials up to eleven. We tested whether sequence length influenced Go response amplitudes, hypothesizing that longer sequences, representing greater unexpected contextual novelty, would lead to enhanced novelty responses.

A computational model based on inhibitory plasticity [57] predicted that novelty responses increase with the number of preceding sequence repetitions, before reaching saturation. Interestingly, our results support this prediction: on Day 4, Go responses increased with the number of preceding images. Notably, we did not observe a clear saturation effect, likely due to the limited sequence range relative to theoretical models. This pattern was present in EXC and VIP cells but absent in SST cells (Figures 1O-1Q). On the remaining days, the trend persisted only in EXC cells, consistent with the reduced VIP responses to the Go outside Day 4 (Figures S2A-S2F). Repeating the analysis while restricting it to common cells tracked across all six experimental days yielded similar results (Figures S2G-S2I), confirming the robustness of the effect.

### Cell type-specific responses to natural images

We next examined how excitatory and inhibitory neuron populations respond to natural image stimuli. While excitatory responses to natural stimuli have been well characterized [51,58–62], inhibitory interneuron responses, particularly those of SST and VIP cells, remain largely unexplored. Prior work has largely focused on their responses to artificial stimuli such as gratings [63], rather than complex, naturalistic inputs [62]. Recent findings show that visual cortex responses differ markedly between artificial and natural stimuli [48], underscoring the need to study inhibitory activity under naturalistic conditions.

To study these responses, we restricted our dataset to neurons recorded continuously across all six days, enabling within-cell comparisons over time while minimizing variability due to fluctuations in the recorded populations. This subset included 694 excitatory,53 SST and 177 VIP cells. Analyses repeated on this subset of common cells reproduced the results shown in Figure 1, confirming consistency across populations (Figure S1).

To systematically quantify image responsiveness, we applied uniform, conservative criteria to classify cells (see Methods), following manual data validation. A neuron was considered responsive to a specific image if its activity during repeated presentations of that image significantly exceeded baseline activity measured during the pre-task grey screen presentation, in a substantial proportion of trials. Image responsiveness varied across neurons, with some showing consistent responses across trials and days, while others were more variable (Figures 2A-2B).

**Fig 2.**
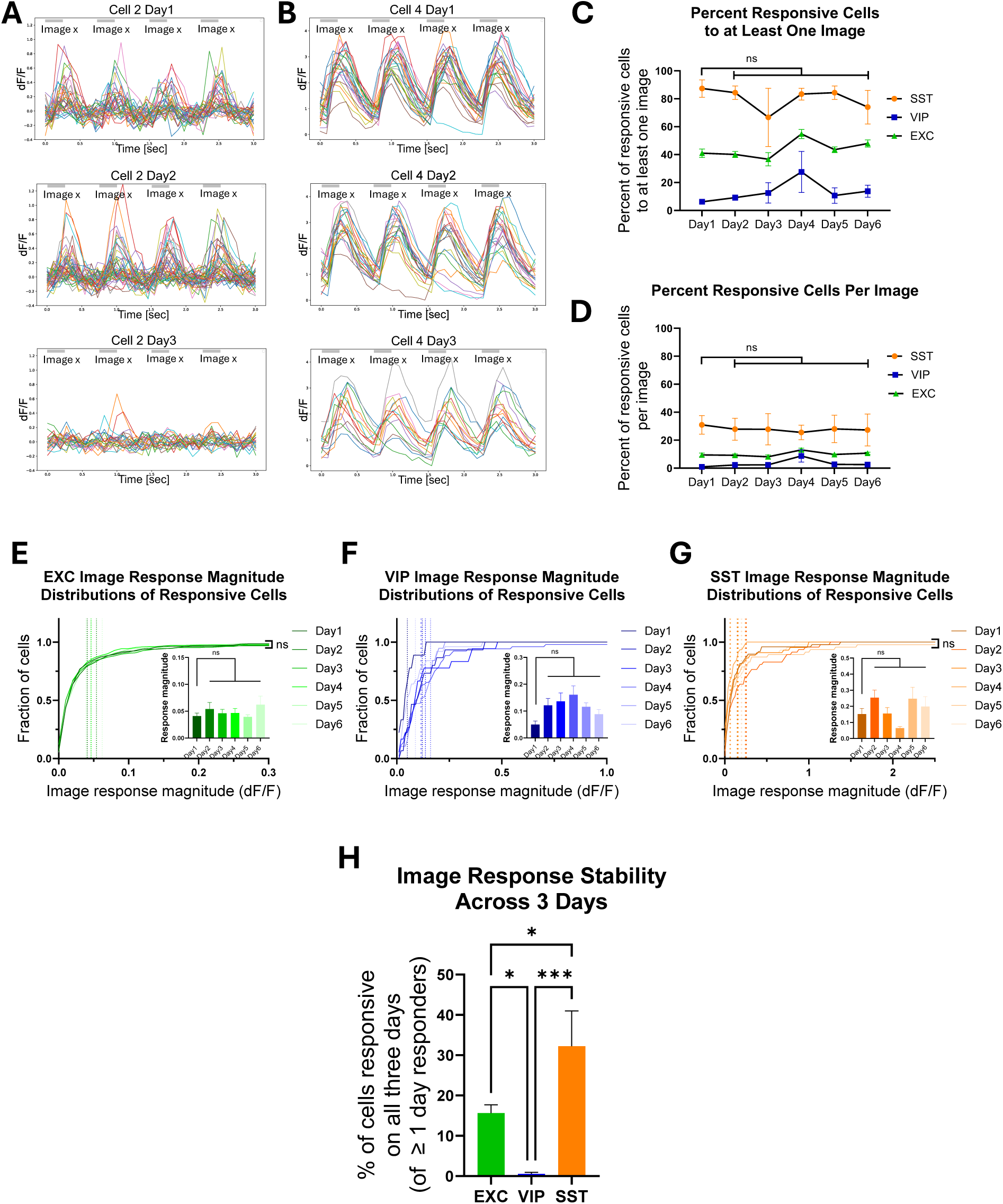
Cell type-specific stability of neuronal responses to natural images. (A-B) Example neuronal responses to repeated presentations of the same image across days (each colored trace represents one trial with four image repetitions). (A) EXC neuron: example neuron that was responsive to a specific image on Days 1-2 but lost responsiveness on Day 3. (B) SST neuron: example neuron showing stable responses to a specific image across Days 1-3. (C) The percentage of neurons responsive to at least one image per day, out of the total number of neurons recorded on all days, remained stable across days. On Day 1, SST neurons showed the highest proportion of responsive cells, and VIP the lowest (two-way RM ANOVA; cell type F(2, 10) = 47.1, p < 0.0001; day F(5, 50) = 3.5, p = 0.008; Tukey’s post hoc comparisons on Day 1 showed significant differences between all cell types, p < 0.001. No significant differences were observed between days when compared to Day 1). (D) Percentage of image responsive cells per image, averaged across the eight images in each set. EXC and SST populations showed stable responsiveness across days, whereas VIP neurons exhibited a significant increase on Day 4 (two-way RM ANOVA; cell type F(2, 10) = 11.6, p = 0.002; day F(5, 50) = 1.00, p = 0.4. Tukey’s post hoc on Day 1 showed significant differences between SST vs. VIP and EXC, p < 0.0001. No significant differences were observed between days when compared to Day 1, except VIP Day 1 vs. Day 4). (E-G) Distributions of image response magnitudes for neurons classified as image responsive on that day were stable at the population-level. (E) EXC, (F) VIP, and (G) SST populations all showed stable response distributions across days (Kolmogorov-Smirnov tests comparing Day 1 vs. Days 2-6, Bonferroni-adjusted p > 0.2 for all comparisons). Inset: average response magnitudes per day also did not differ significantly (one-way ANOVA; EXC, F(5,1859) = 0.7, p = 0.6; VIP, F(5,132) = 0.8, p = 0.5; SST, F(5,271) = 2.0, p = 0.07; Dunnett’s post hoc tests, all adjusted p > 0.1). (H) Percent of cells stably responsive to the same image on three consecutive days out of the total number of responsive cells to that image across those days, calculated per mouse for Days 1-3 and Days 4-6 and averaged (EXC, n = 6 mice; VIP, n = 4; SST, n = 3). SST cells had significantly higher stability at the single-cell level (one-way ANOVA; F(2, 10) = 14.5, p = 0.001 followed by Tukey’s post hoc: SST vs VIP, p = 0.0008; SST vs EXC, p = 0.03; EXC vs VIP, p = 0.03). Error bars: (C-D,H) ± SEM across mice;(E-G) ± SEM across neurons. ns p>0.05, *p < 0.05, **p < 0.01, ***p< 0.001, ****p<0.0001.

We quantified the percentage of neurons responsive to at least one image (Figure2C), and the percentage of responsive neurons per image (Figure2D). Excitatory response patterns aligned with prior reports [61,15]. SST neurons were the most responsive (87% ± 6 on Day 1), followed by EXC (41% ± 3), and VIP cells (6% ± 2). Notably, this characterization of natural image responses is among the first to be conducted separately for SST and VIP inhibitory cells. Importantly, the proportion of image responsive cells remained largely stable across days for each cell type, with the exception of an increase in VIP responsiveness on Day 4 (Figure 2D).

### SST cells exhibit the highest single-cell stability of image responses across days

Having established that the number of image responsive cells remained stable across days—except for VIP cells on Day 4 (Figure2D) —we next examined the stability of response magnitude at the population level. For each day, we calculated the average and distribution of response magnitudes among neurons classified as image responsive on that day. Both measures were consistent across days for each cell type (Figures 2E-2G, Figures S4B-S4C), indicating that the population-level response of responsive neurons was stable over time.

We then asked whether this stable population response reflected stable responses from the same neurons. To assess single-cell stability, we quantified, for each image, the proportion of neurons that were responsive to that image on all three consecutive days, relative to those that were responsive to that image on at least one of those days. These proportions were first averaged across the eight images within each set, and then across the familiar (Days 1-3) and novel (Days 4-6) sets, yielding a single stability value per mouse.

Despite population-level stability, a large fraction of image responsive cells did not remain responsive across days (Figure 2H). Notably, SST cells exhibited the highest stability, with 32% ± 9 of responsive cells consistently responsive to the same image across three days, approximately twice that of EXC cells (16% ± 2). VIP cells showed minimal stability. Importantly, stability for both EXC and SST populations was significantly above chance based on a shuffle-based null model (exceeded the 99th percentile of the null distribution). In addition, as expected, response stability was higher when comparing two-days responsiveness to three-days (FigureS4D). Together, these results show that while stable population-level responses arise from a mix of stable and unstable cells, SST neurons uniquely maintain the most stable image responses over time.

### SST cells exhibit the highest single-cell stability of image change (Go) responses across days

We next assessed the stability of neuronal responses to image changes (Go responses). Neurons were classified as Go responsive if their response to an image change exceeded their response to the preceding image (see Methods). As shown in Figure3C, SST and EXC cells exhibited the highest percentage of Go responsive cells, followed by VIP cells. The introduction of a novel image set on Day 4 led to an increase in Go responsive EXC cells. At the population level, both the proportion of Go responsive neurons and the average and distribution of their response magnitudes were stable across days for EXC and SST cells (Figures 3D-3F), indicating consistent population-level Go responses over time. VIP cell responses were largely stable across days, however no Go responses were detected on Day 2.

**Fig 3.**
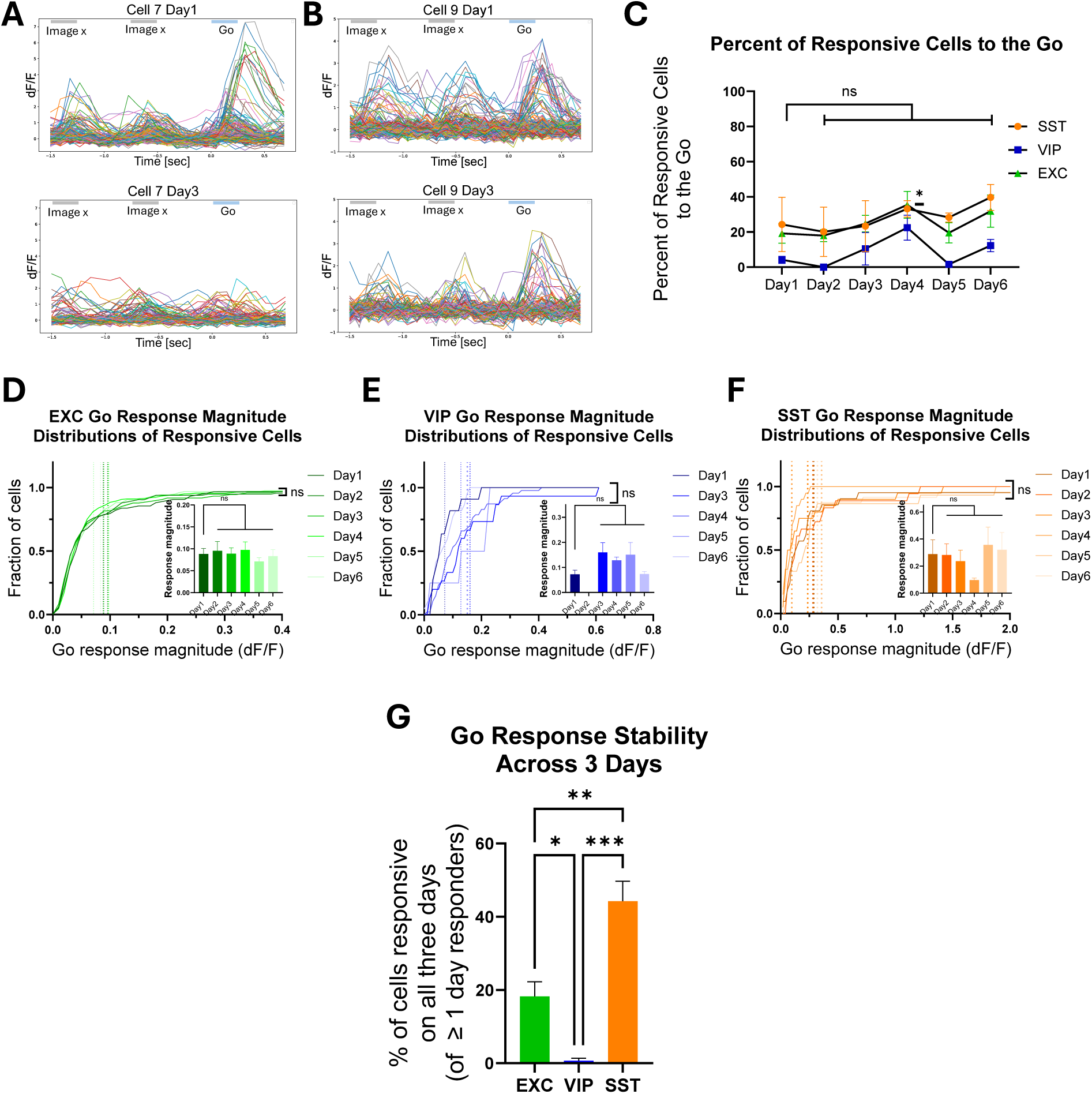
Cell type-specific stability of image change (Go) responses. (A-B) Example neuronal responses to image change (Go) across days (each colored trace represents a trial showing the last two image repetitions followed by an image change). (A) EXC neuron: example neuron that was responsive to the Go on Day 1 but not on Days 2-3. (B) SST neuron: example neuron with stable Go responses on Days 1-3. (C) The percentage of cells responsive to the Go per day, out of the total number of neurons recorded on all days, remained stable over time. Percentages were stable across days except for a significant increase in EXC neurons on Day 4 (two-way RM ANOVA; day F(5, 50) = 7.1, p < 0.0001. Tukey’s multiple comparisons between days were ns, except for Day 4 vs. Day 1 in EXC, p = 0.04). (D-F) Distributions of Go response magnitudes for neurons classified as Go responsive on that day were stable at the population level. (D) EXC neurons showed stable distributions of Go responses among neurons classified as Go responsive on each respective day (Kolmogorov-Smirnov tests, Bonferroni-adjusted p > 0.9). Inset: average response magnitudes also showed no significant differences (one-way ANOVA; F(5, 907) = 0.3, p = 0.9; Dunnett’s post hoc, p > 0.9). (E) VIP responses were stable on Days 1,3-6 (Bonferroni-adjusted p > 0.3); no Go responses were detected on Day 2. Inset: average responses on Days 3-6 were not significantly different from Day 1 (ANOVA; F(4, 94) = 3, p = 0.03; Dunnett’s, p > 0.05). (F) SST neurons showed stable response distributions and averages across days (Bonferroni-adjusted p > 0.3; ANOVA; F(5, 112) = 0.7, p = 0.7; Dunnett’s, p > 0.5). (G) The Percent of cells stably responsive to the Go on three consecutive days out of the total Go responsive cells across those days, calculated per mouse for Days 1-3 and Days 4-6 and averaged. SST neurons showed significantly higher stability at the single-cell level (one-way ANOVA F(2, 10) = 24, p = 0.0001, followed by Tukey’s multiple comparisons test: SST vs VIP, p = 0.0001; SST vs EXC, p = 0.003; EXC vs VIP, p = 0.02). Error bars: (C,G) ± SEM across mice; (D-F) ± SEM across neurons. ns p>0.05, *p < 0.05, **p < 0.01, ***p< 0.001, ****p<0.0001.

To assess single-cell stability, we calculated the proportion of cells that were Go responsive on all three days relative to those responsive on at least one day, computed separately for Days 1-3 and Days 4-6, and then averaged to obtain a single value per mouse. As with image responses, a substantial fraction of cells across all cell types exhibited unstable responses. Notably, SST cells exhibited the highest single-cell stability, approximately twice that of EXC cells (SST: 44% ± 5, EXC: 18% ± 4, Figure3G), while VIP cells displayed minimal stability. Stability in both EXC and SST populations significantly exceeded chance levels based on a shuffle-based null model (exceeded the 99th percentile of a null distribution). As expected, stability was higher across two-day comparisons than three-day comparisons (Figure S4E).

### VIP and EXC cells exhibit comparable single-cell stability of omission responses across days

To complete our analysis of response stability across all stimulus types, we examined neuronal responses to image omissions. In contrast to other stimulus types, omission responsiveness was highest among VIP neurons and lowest among SST cells (VIP: 60% ± 13, EXC: 7% ± 2, SST: 6% ± 0, Figure4C). While the population-averaged excitatory responses to omissions suggested a lack of activation (Figure1I), a subset of EXC cells exhibited increased activity during omissions and were classified as omission responsive, revealing heterogeneity masked in the population average. Due to the low number of omission responsive SST cells, stability analysis was restricted to EXC and VIP populations. At the population level, both the fraction of omission responsive cells (except VIP on Day 4) and their average and distribution of response magnitudes were stable across days (Figures 4D-4E), indicating consistent omission evoked population responses over time. Stability at the single-cell level, calculated as described in the preceding subsections, was comparable between these EXC and VIP cells, with both exhibiting a high proportion of unstable neurons (Figure4F). Nonetheless, their stability exceeded chance levels based on a shuffle-based null model. To further contextualize these findings, we compared the stability of neuronal responses across all tested conditions: image presentations, image changes, and omissions (Figures 4G-4I).

**Fig 4.**
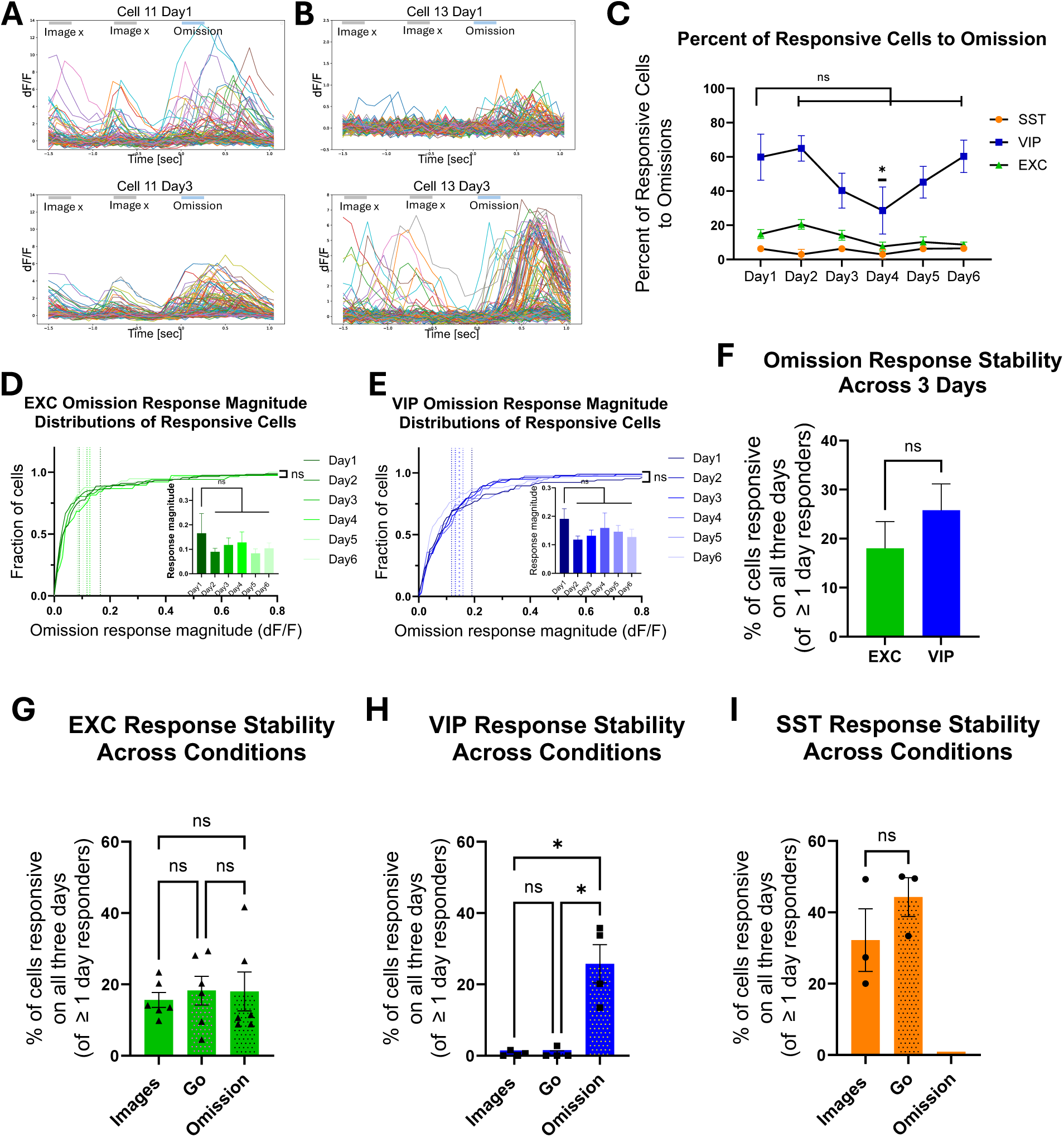
Cell type-specific stability of omission responses. (A-B) Example neuronal responses to image omissions across days (each trace represents a single trial showing two image presentations followed by an omission). (A) EXC neuron: example neuron with stable omission responses on Days 1-3. (B) VIP neurons: example neuron with stable omission responses on Days 1-3. (C) The percentage of cells responsive to omission per day, out of the total number of neurons recorded on all days, remained stable over time, except for VIP Day 4 (two-way RM ANOVA; day F(5,45)=1.9, p=0.1, Tukey’s post hoc in VIP Day 4 vs. Day 1, p=0.02; all other comparisons are ns). (D-E) Distributions of omission response magnitudes for neurons classified as omission responsive on that day were stable at the population level. (D) EXC neurons showed consistent omission response distributions across days (Kolmogorov-Smirnov tests vs. Day 1, Bonferroni-adjusted p > 0.9). Inset: average response magnitudes per day were also not significantly different (one-way ANOVA F(5, 437) = 0.5, p = 0.8; Dunnett’s post hoc, p > 0.5). (E) VIP omission response distributions and means were stable across days (Bonferroni-adjusted p > 0.9; ANOVA F(5, 504) = 1, p = 0.4; Dunnett’s post hoc, p > 0.1). (F) The percent of cells stably responsive to omission on three consecutive days out of the total omission responsive cells across those days, calculated per mouse for Days 1-3 and Days 4-6 and averaged. EXC and VIP neurons showed comparable stability at the single-cell level (unpaired t-test, t(8) = 0.9, p = 0.4). (G-I) Comparison of the percentage of stably responsive cells across the three stimulus conditions: images, Go, and omissions. (G) EXC neurons were similarly stable across all three conditions (RM one-way ANOVA F(1.42, 7.1) = 0.1, p = 0.8, Tukey’s multiple comparisons were ns). (H) VIP neurons were only stably responsive to omissions (F(1, 3) = 19.4, p = 0.02 Tukey’s: image vs. Go, ns; image vs. omission vs. image and Go, p = 0.04). (I) SST neurons showed similar stability for image and Go responses (paired t-test, t(2) = 1.3, p = 0.3). Error bars: (C,F-I) ± SEM across mice; (D-E) ± SEM across neurons. ns p>0.05, *p < 0.05, **p < 0.01, ***p< 0.001, ****p<0.0001.

To further assess population-level stability, we performed logistic regression decoding of image changes and omissions from the activity of all individual neurons, training and testing the model separately for each day on the population activity. All cell types exhibited robust decoding accuracy within days, with VIP decoding performance for Go responses increasing on Day 4 (Figures S5). These findings further support the conclusion that population-level novelty responses remain stable across days, even as the identity of responsive neurons changes.

In summary, these results reveal a dissociation between population-level and single-cell stability. Across conditions, the responsive neuronal population maintains comparable size and response magnitude across days, however, the identity of responsive cells varies from day to day. Notably, SST neurons consistently displayed the highest single-cell stability, suggesting a unique role in maintaining stable stimulus representations. EXC neurons showed consistent stability across conditions, indicating that single-cell response stability is preserved within the excitatory population across the specific conditions tested in this experiment. VIP neurons displayed stable responses only in the omission condition, pointing to a selective stabilization in response to negative prediction errors. This dissociation points to a more nuanced and condition-dependent role for VIP neurons in novelty processing than previously appreciated. Together, these results represent a novel dissociation in SST, EXC, and VIP neuron stability, reflecting distinct roles in sensory processing.

### Go and omission responses engage largely non-overlapping neurons

In addition to stability, a key feature of novelty responses is the nature of the information they encode [52–56, 64]. We first asked whether individual neurons respond to multiple forms of novelty—namely, Go and image omissions. We then examined whether Go responses carry general or stimulus-specific information.

We first quantified the proportion of cells responsive to at least one condition across days (Figure5A). The fraction of responsive neurons remained stable across days for all cell types. To examine novelty response overlap, we quantified the proportion of cells responsive to each task condition and measured how these populations intersected, averaging the proportions across Days 1-6. For completeness, we also report the overlap between Go and image responsive cells, which will be further addressed in the next subsection. In the SST population, nearly all Go responsive cells also responded to images, indicating substantial overlap between these response types. However, there was no overlap between Go and omission responsive cells (Figures 5D,5G). In the EXC population, approximately 60% of Go responsive neurons were also image responsive, while only a small fraction (4%) were responsive to both Go and omission conditions (Figures 5B,5E). VIP cells showed low overlap across all response types, including a generally low intersection between Go and image responses, with a slight increase on Day 4 (Figures 5C, 5F).

**Fig 5.**
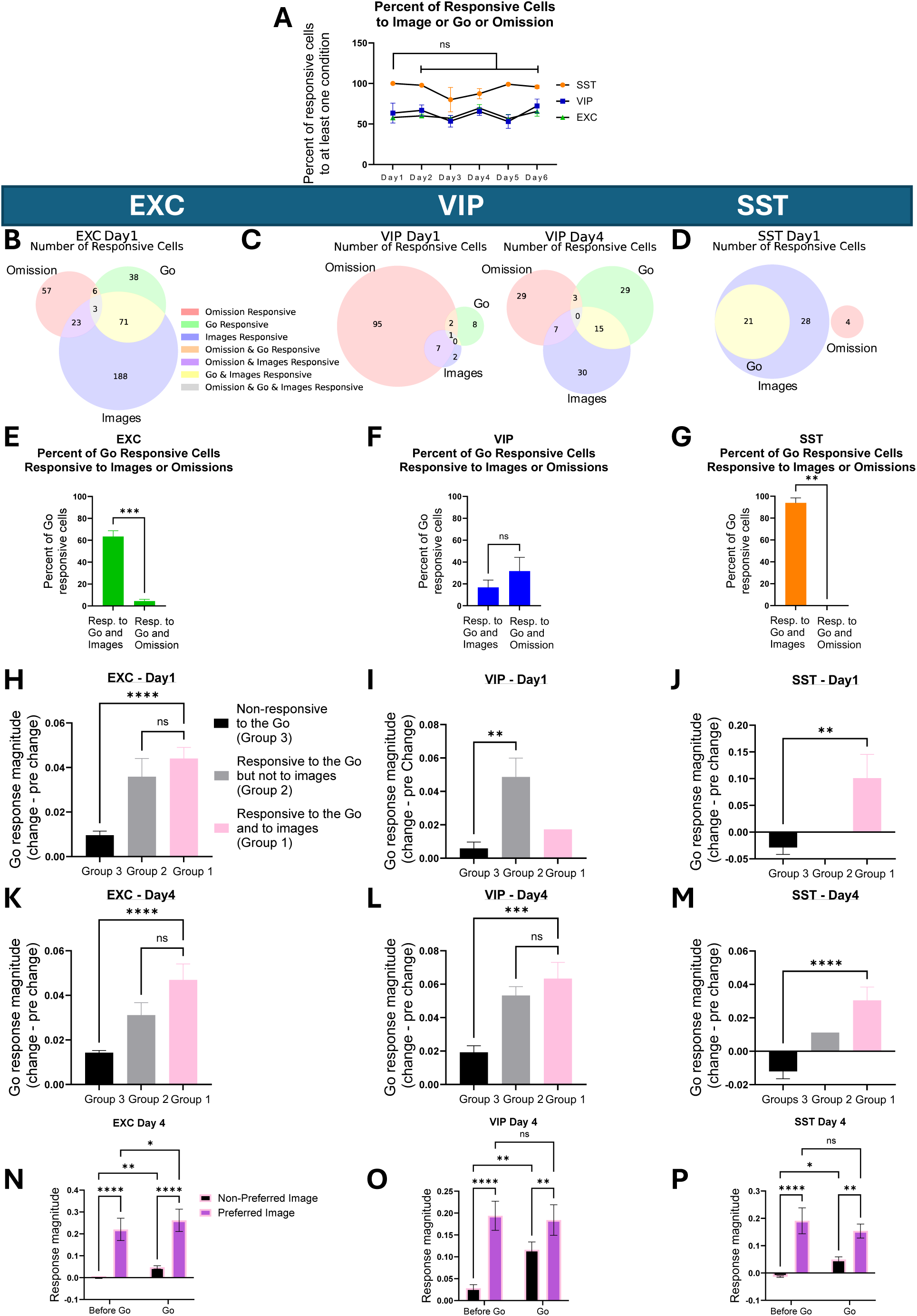
Go responses include both stimulus-specific and non-specific components across cell types, with VIP neurons shifting toward stimulus-specific responses on Day 4. (A) Percentage of neurons responsive to at least one condition (images, Go, omissions) remained stable across days for all cell types (two-way RM ANOVA; day F(5,50) = 1.1, p = 0.4, Tukey’s post hoc showed no significant differences). (B-D) Overlap between image, Go, and omission responsive cells, each circle represents the number of neurons responsive to images, Go or omissions on a given day, pooled across all mice. Day 1 is shown for EXC and SST; both Day1 (left) and Day 4 (right) shown for VIP. (E-G) Among Go responsive neurons, a higher proportion were also image responsive than omission responsive in EXC and SST cells (paired t-test; EXC: t(5)=10.9, p<0.001; SST: t(2)=20.5, p<0.01; VIP: t(3)=1.3, p=0.3). The proportions shown represent averages across Days 1-6. (H-J) Go response magnitude (difference between image change and preceding image responses) on Day 1 for three groups of neurons: “*Image + Go Responsive*” (Group 1-pink: n=74 EXC cells, 1 VIP, 32 SST), “*Go-Only Responsive*” (Group 2-gray: n=44 EXC, 10 VIP), and “*Go-Nonresponsive*” (Group 3-black: n=576 EXC, 166 VIP, 21 SST), (EXC: one-way ANOVA F(2, 691) = 25, p < 0.0001, Dunnett’s multiple comparisons: responsive to Go and Images vs. non-responsive to Go, p < 0.0001; VIP: unpaired t-test t(174) = 3, p = 0.007; SST: t(51) = 3, p = 0.002). (K-M) Same analysis on Day 4. Top: Go response magnitude on Day 4 for the same three groups. “*Image + Go Responsive*” (pink: n=170 EXC cells, 15 VIP, 19 SST), “*Go-Only Responsive*” (gray: n=56 EXC, 32 VIP, 1 SST), and “*Go-Nonresponsive*” (black: n=468 EXC, 130 VIP, 33 SST), (one-way ANOVA; EXC: F(2, 691) = 26, p < 0.0001, Dunnett’s multiple comparisons: responsive to Go and Images vs. non-responsive to Go, p < 0.0001; VIP: F(2, 174) = 14, p < 0.0001, post hoc: responsive to Go and Images vs. non-responsive to Go, p = 0.0003; SST: unpaired t-test, t(50) = 5, p < 0.0001). (N-P) Responses of the *Image + Go Responsive* group to preferred vs. non-preferred images, before and during the Go stimulus. Responses were significantly higher for preferred vs. non-preferred images, both before and during the Go, and increased from before Go to Go (two-way ANOVA; EXC: time: F(1,169) = 17, p < 0.0001; image preference F(1,169) = 20, p < 0.0001, Fisher’s LSD post hoc; group difference before Go p < 0.0001, Go p < 0.0001; within non-preferred image group before vs. Go p = 0.004; within preferred image group before vs. Go p = 0.01; VIP: time: F(1,14) = 7, p =0.02; image preference F(1,14) = 21, p = 0.0004, Multiple comparisons-group difference before Go p < 0.0001, Go p = 0.008; within non-preferred image group before vs. Go p = 0.001; within preferred image group before vs. Go p = 0.7; SST: time: F(1,18) = 0.9, p =0.3; image preference F(1,18) = 25, p < 0.0001, Multiple comparisons- group difference before Go p < 0.0001, Go p = 0.001; within non-preferred image group before vs. Go p = 0.03; within preferred image group before vs. Go p = 0.2). Error bars: (A, E-G) ± SEM across mice; (H-P) ± SEM across neurons. ns p>0.05, *p < 0.05, **p < 0.01, ***p< 0.001, ****p<0.0001.

These results reveal distinct patterns of neuronal recruitment across task conditions: while image and Go responses overlap substantially in the EXC and SST populations, Go and omission responses engage largely separate populations across all cell types. This dissociation may reflect differential roles in encoding positive versus negative prediction errors [64].

### Go responses include both stimulus-specific and non-specific components across cell types

An important question is what information is encoded by the cells responding to the Go stimulus—specifically, whether the contextual novelty response is a broad response, namely non-specific to the image stimulus during the Go, or is it a stimulus-specific representation of the image identity during the Go [22,23, 52–56]. Furthermore, whether the information content remains consistent across days or is modulated by the absolute novelty condition introduced on Day 4. Namely, does the neuronal activity encode both the occurrence of novelty—image change and the content of the novel stimuli—the image presented, giving the brain the ability to detect the unexpected and understand what it is.

To address this, we categorized neurons into three groups: (1) “*Image + Go Responsive*”: neurons responsive to both Go stimuli and images, (2) “*Go-Only Responsive*”: neurons responsive only to Go stimuli, and (3) “*Go-Nonresponsive*”: neurons not classified as Go-responsive.

In the excitatory population on Day 1, both “*Image + Go Responsive”* and “*Go-Only Responsive”* neurons showed significantly elevated responses during the Go compared to the preceding image (Figure5H). Neurons in group 3, exhibited a mild increase in activity, indicating that novelty-related signals are present across the excitatory network. The same pattern was observed on Day 4 (Figure5K).

In the SST population, on Day 1, no “*Go-Only Responsive”* cells were detected. Although the overall SST population showed no significant Go response (Figure1I), neurons in the “*Image + Go Responsive”* group exhibited a significant activity increase, consistent with the SST Go responsive cells shown in Figure 3. Conversely, “*Go-Nonresponsive*” neurons showed a significant decrease in activity. This pattern persisted on Day 4 but was reduced, consistent with the population-level trends shown in Figure1 (Figures 5J,5M).

In contrast, the VIP population exhibited markedly different response dynamics. On Day 1, Go responses were exclusively driven by “*Go-Only Responsive”* neurons. On Day 4, however, all three groups contributed to the Go responses, with groups “*Image + Go Responsive”* and “*Go-Only Responsive”* contributing the most (Figures 5I,5L).

The response in the *Go-Only Responsive* group reflects a novelty response that is not stimulus specific. To determine whether the response in the *Image + Go Responsive* group is stimulus-specific, we examined neuronal responses to preferred and non-preferred images both before and during the Go stimulus. Preferred and non-preferred images were defined for each neuron based on responses to repeated image presentations prior to the Go (as in Figure2). Preferred images evoked stronger responses both before and during the Go. This finding, in addition to the increased response during the Go, indicates that, based on neuronal activity, both the occurrence of an image change and the identity of the presented image are encoded, reflecting a stimulus-specific novelty response (Figures 5N-5P).

To summarize, EXC cells exhibited stimulus-specific novelty response, and also a non-specific stimulus response, on both Day 1 and Day 4. SST cells demonstrated stimulus-specific novelty responses across both days. Notably, VIP cells transitioned from a non-specific stimulus response on Day 1 to exhibiting both specific and non-specific responses on Day 4. Our results corroborate the findings of Bastos et al. [23] and Furutachi et al. [22] regarding stimulus-specific responses among EXC neurons, but extends them by identifying non-specific responses in both EXC and VIP neurons, with the presence of the non-specific response in EXC neurons being of particular interest. Finally, we observed the emergence of stimulus-specific responses in VIP neurons on Day 4, representing a key new finding.

Together, these findings suggest that novelty responses to the Go comprise both a non-specific novelty signal and an enhanced representation of the unexpected visual stimulus itself. This dual mechanism may amplify the representation of novel input features, thereby facilitating learning and adaptive processing. The observed change in information content within the VIP population on Day 4 indicates that encoding is dynamic and modulated by absolute novelty. Moreover, the differential responses across cell types imply distinct functional roles in novelty detection: EXC and VIP neurons integrate both nonspecific and stimulus-specific novelty signals, whereas SST neurons selectively amplify the representation of stimulus-specific novel stimuli.

### Electrophysiological recordings reveal novelty-dependent changes in VIP dynamics

Lastly, we hypothesized that the change in the properties of the responses of the VIP cells on Day 4, might also be manifested in the firing dynamics. For instance, maybe there is only a change in the magnitude of the activity or maybe the dynamics in addition to the magnitude are changed.

To test this, we analyzed Neuropixels recordings from a parallel dataset collected by the Allen Institute, offering high temporal resolution. This experiment included two recording days—familiar (Day 1) and novel (Day 2)—with both active and passive conditions. Two images were repeated across both days (‘shared’), while six were unique to each day (‘non-shared’), allowing assessment of an additional facet of novelty, namely absolute novelty specificity (Figure 6A). Acute recordings prevented tracking the same cells across days, but enabled precise examination of firing patterns. To identify cell types, two genetic mouse lines were used expressing channel rhodopsin in SST+ and VIP+ neurons. Optotagging enabled classification of VIP and SST neurons, while spike waveform properties were used to distinguish putative excitatory and parvalbumin (PV) neurons [65–67]. Recordings from VISp and VISl regions yielded EXC (3842 ± 317 neurons, *n*=27 mice), PV (759 ± 105, *n*=27), SST (220 ± 26, *n*=19), and VIP neurons (17 ± 5, *n*=7).

**Fig 6.**
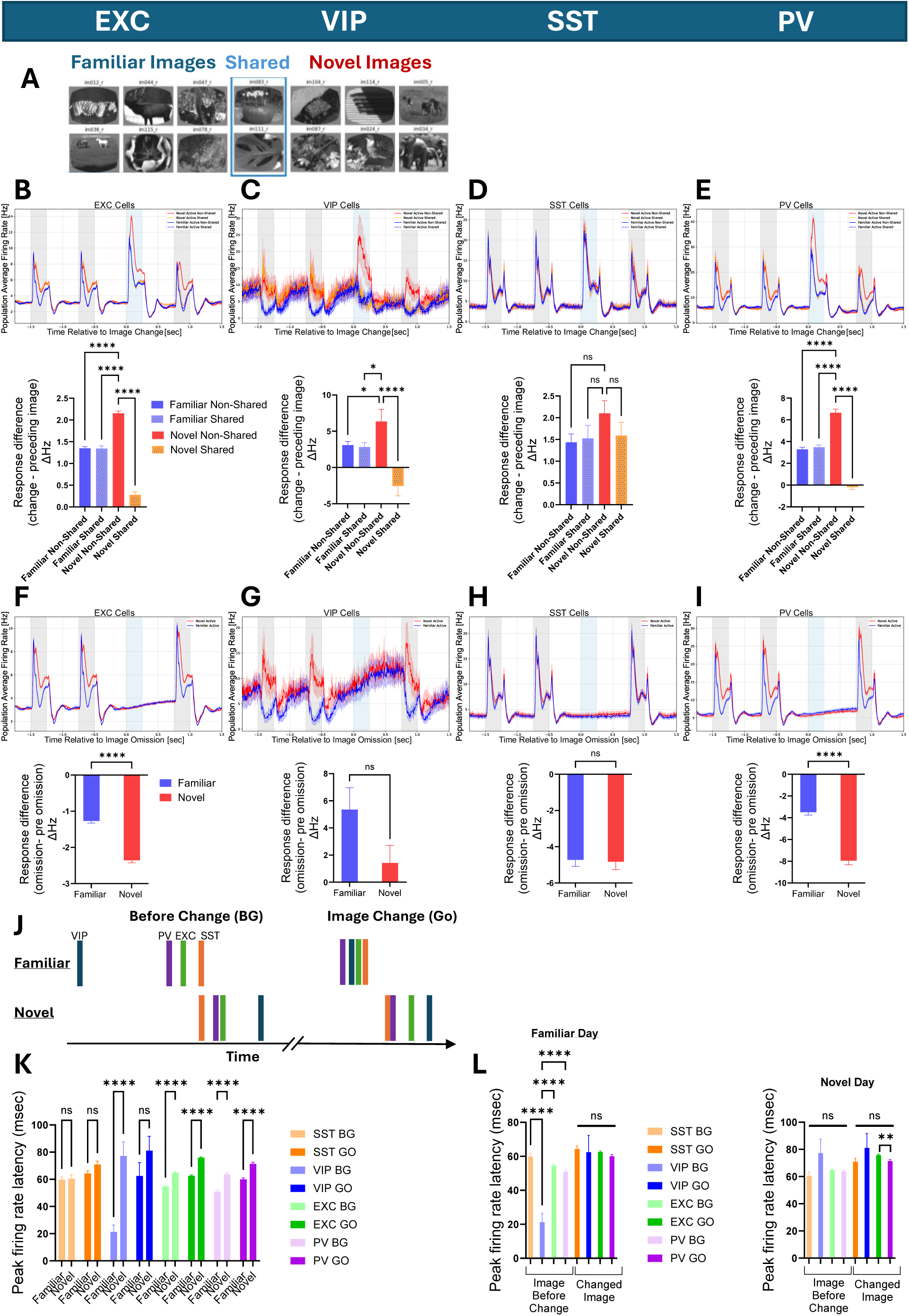
Electrophysiology uncovers distinct VIP firing rate patterns during familiar and novel conditions. (A) Mice performed a visual change detection task in which natural images were presented repeatedly until an image change occurred. 5% of non-change trials included stimulus omissions. On Day 1 (familiar day), mice were shown a set of familiar images. On Day 2 (novel day), a new image set was introduced, comprising six novel images (“non-shared”) and two familiar images (“shared”). Neuronal activity was recorded acutely each day using Neuropixels probes. (B-E) Top: population-averaged responses to image presentations (gray) and image changes (Go, blue) across familiar and novel days of EXC, PV, SST, and VIP cells identified by optotagging (EXC: 3842 ± 317 neurons from n=27 mice, SST: 220 ± 26 neurons from n=19 mice, VIP: 17 ± 5 neurons from n=7 mice, PV: 759 ± 105 neurons from n=27 mice). Bottom: on the novel day, responses to non-shared images during the Go were enhanced in EXC, PV, and VIP (one-way ANOVA followed by Dunnett’s post hoc comparing all days to novel non-shared images; EXC: F(3, 15366) =172, p < 0.0001, All multiple comparisons were significant, p < 0.0001; VIP: F(3, 64) =12, p < 0.0001, multiple comparisons, p < 0.05; SST: F(3, 878) =1, p =0.3, multiple comparisons were ns; PV: F(3, 3032) =120, p < 0.0001, multiple comparisons, p < 0.0001). (F-I) Top: population-averaged responses to stimulus omissions (blue) across days. Bottom: comparisons between the days (unpaired t-test; EXC: t(7683) = 12, p < 0.0001; VIP: t(32) = 2, p=0.1; SST: t(439) = 0.2, p=0.8; PV: t(1516) = 10, p < 0.0001). (J) Schematic illustration of peak response latencies for each cell type during image presentation before the Go (BG) and during the Go (Go), for both days. (K) Response latency increased on the novel day in EXC and PV cells for both BG and Go images, and in VIP cells during BG (two-way ANOVA; day, F(1, 7797) = 70, p<0.0001, Sidak’s post hoc between days were significant for EXC and PV and VIP during BG, p<0.0001). (L) Left: on the familiar day, latencies differed between cell types during BG but converged during the Go. Right: on the novel day, latencies were comparable across cell types for both BG and Go. VIP neurons showed the largest shift, from earliest on Day 1 to latest on Day 2 (Familiar: one-way ANOVA for the Go F(3, 2267) =2, p =0.1, one-way ANOVA for the BG F(3, 1547) =14, p<0.0001 followed by Tukey’s post hoc. Novel: Go F(3, 2316) =4, p =0.005 followed by Tukey’s post hoc test only EXC vs PV significant, BG F(3, 1667) =1, p=0.2). Error bars: ±SEM across neurons. ns p>0.05, *p < 0.05, **p < 0.01, ***p< 0.001, ****p<0.0001.

We observed population-level activity patterns similar to those seen in the calcium imaging dataset: EXC and VIP cells showed increased response to the changed image compared to the preceding image on both days (Figures 6B-6E), and VIP neurons exhibited ramping activity during omissions (Figures 6F-6I). The responses of PV neurons highly resembled that of excitatory cells. One exception was the absence of a strong SST suppression on the novel day, potentially due to differences in cell selection bias or spike-burst dynamics across recording methods [68], this should be systematically assessed in future experiments.

Critically, the fine temporal resolution of Neuropixels revealed changes in the firing dynamics induced by novelty. On the familiar day, VIP neurons showed ramping responses between images and suppression at image onset followed by ramping activity until image offset. Upon image change, the suppression was reduced. Interestingly, On the novel day, this pattern reversed— image onset triggered strong VIP activation and upon image change the activity increased and remained high. In summary, VIP neurons reversed from being inhibited by images stimuli to being active upon image presentation (Figure 6C).

The presence of shared and non-shared images enabled testing for absolute novelty specificity. Upon image change EXC, VIP, and PV neurons showed enhanced responses to the non-shared (novel) images on Day 2, while responses to shared images resembled those on the familiar day (Figures 6B-6E, S6A-S6D). Excitingly, this highlights that the absolute novelty signal is highly specific and is not uniform across the novel day, adding another aspect to our finding of stimulus-specific contextual novelty responses of these cells.

Next, we examined the response latency by evaluating the time to peak response in response to the image before the change (BG) and the image following the change (Go) [69–72]. Two main phenomena emerged. First, response latencies increased on the novel day in EXC neurons, consistent with prior reports [73,74]. A similar increase was observed in PV neurons, while VIP neurons showed delayed responses specifically during the BG image. SST latencies remained unchanged (Figure6K). Secondly, the relative order of peak firing latencies across cell types shifted. On the familiar day, during BG, VIP neurons responded earliest, followed by PV and EXC, with SST firing latest. During the Go, all cell types exhibited similar latencies. However, on the novel day, latencies were comparable across cell types during both BG and Go (Figures 6J,6L). Notably, VIP neurons exhibited the most pronounced shift: from being the earliest to reach peak activity on the familiar day to the latest on the novel day. Furthermore, when comparing responses to shared and non-shared, latency during Go did not increase relative to BG in EXC and PV neurons, and in VIP neurons latency even decreased slightly, when the Go image was shared (FigureS6J). This further supports the stimulus-specific nature of latency changes.

Together, these findings confirm and extend our calcium imaging results. They highlight that VIP cells undergo the strongest modulation on the novel day, showing changes in firing pattern, stimulus specificity, and temporal dynamics. These changes in activity patterns may be linked to the changes in information content encoded by these cells that were observed in the calcium imaging data under conditions of absolute novelty.

## Discussion

Detecting and processing novelty is critical for learning and survival, yet how such novelty is represented across neuronal populations—particularly at the level of single cells and across cell types—remains unresolved. Are the responses of individual EXC, SST, and VIP neurons to different forms of novelty stable over time, or do stable population-level representations emerge despite single-cell variability? Do neurons that respond to contextual novelty carry fixed information across days, or can their coding properties be dynamically reshaped?

Here, we show that these subtypes differ in their responses to visual input and novelty, suggesting distinct roles in cortical processing. To our knowledge, this is the first systematic assessment of response stability across days in excitatory and inhibitory neurons under novelty conditions. All cell types exhibited both stable and unstable responsive neurons, yet SST cells showed the highest proportion of stably responsive cells, followed by EXC and VIP cells. Moreover, VIP neurons displayed stable responses only to omissions. These findings raise important questions about the relationship between stability at the level of individual neurons and at the population level. To further understand the nature of novelty representations, we examined the information content of contextual novelty responses. We found distinct patterns of contextual novelty encoding across cell types: EXC cells encoded both stimulus-specific and non-specific novelty, VIP neurons transitioned from non-specific to both stimulus-specific and non-specific encoding under absolute novelty conditions, revealing a flexible and previously unrecognized coding strategy. Summarizing our findings and current knowledge in the field, we hypothesize the following roles for SST, EXC, and VIP cells in cortical processing.

### SST interneurons exhibit high response stability and may contribute to familiar stimulus encoding

We found that SST interneurons exhibit stable responses at the population level and greater single-cell stability compared to both EXC and VIP neurons. To our knowledge, this is the first analysis to directly quantify the stability of SST responses to natural visual stimuli using longitudinal calcium imaging. We hypothesize that SST interneurons play a critical role in visual recognition memory by encoding and storing representations of familiar stimuli. Their high response stability may reflect their integration into memory related ensembles and a role in maintaining familiar visual representations over time, consistent with the idea that stable neuronal activity supports memory storage [75,76]. It is plausible that both EXC and SST neurons contribute together to the storage of familiar representations.

Several lines of evidence from other brain regions support this hypothesis. First, computational modeling has demonstrated that inhibitory networks are more effective than excitatory networks in storing memory, yielding robust activity despite excitatory synaptic volatility [77]. While not focused on specific inhibitory subtypes, the study proposed that inhibitory networks have the potential to control the stability of memory patterns for long periods of time, rather than merely exerting nonspecific suppression. Second, studies in the hippocampus showed that SST activation enhances place cell stability [78, 79] and contributes to predictive goal-related memory [80]. In addition, a recent study [81] reported that inhibitory synapses on the basal dendrites of pyramidal neurons —predominantly originating from SST interneurons—were found to exhibit greater structural stability than other inhibitory or excitatory synapses. Moreover, impairment of these cells is linked to memory dysfunction in Alzheimer’s disease model mice [82], together suggesting that SST cells have a stabilizing role in hippocampal memory circuits. Third, in the visual cortex, an experience-dependent increase in SST activity as visual stimuli become familiar was found [24], consistent with the idea that these neurons encode learned sensory input. Additional support comes from studies in the prefrontal cortex showing that SST ensembles are activated during fear learning and recall, where they promote disinhibition of excitatory neurons involved in learning while suppressing those not engaged [83, 84]. This could align with findings in the motor cortex, of reduction in SST axonal boutons onto excitatory cells upon motor training [85]. These findings may suggest that, in the visual cortex, SST neurons may preferentially inhibit excitatory neurons that are not responsive to specific familiar image stimuli while disinhibiting those that are responsive, thereby enhancing the specific response to the stimuli. Combined together, we propose that SST cells participate in visual memory through stable, selective responses to familiar stimuli.

Moreover, SST interneurons have also been implicated in short-term contextual processing and working memory [86], a key component of change detection tasks [87]. The mechanisms supporting long-term stability of Go responses and short-term responses in SST neurons may be functionally linked, though their interaction across timescales is not yet understood.

An important question is what confers the remarkable stability of SST cells—whether it arises from intrinsic cellular properties or from their connectivity and inputs. Furthermore, it remains to be determined whether stable neurons across different types (EXC, SST, VIP) share common mechanisms that endow them with this stability.

### VIP neurons shift their encoding on the absolute novel day and may support learning

We found that VIP interneurons encoded contextual novelty with a non-specific response on Day 1. This implies that under familiar conditions, VIP neurons signal the presence of novelty without specifying stimulus identity, possibly necessitating other populations to convey that information. By Day 4, when the stimulus set was novel, VIP encoding shifted to include both stimulus-specific and non-specific components. This transition in information content is a novel finding, suggesting an additional role for VIP neurons in novelty representation under absolute novelty conditions. Previous studies have reported only stimulus-specific responses in VIP neurons [22], while other paradigms without reward or training observed no VIP responses to image changes [23]. Our results demonstrate that under trained, task-engaged conditions, VIP cells dynamically alter their encoding profile.

We propose that VIP interneurons play a central role in encoding novel stimuli and serve as key modulators of learning. This role is revealed most clearly on Day 4, as evidenced by elevated VIP activity during both image presentation and the Go stimulus, a shift in firing pattern and temporal dynamics specifically in response to non-shared images, and the emergence of stimulus-specific responses to the Go. As prior studies have linked VIP activity to short-timescale novelty processes [15, 19, 20, 23, 27, 29], recent evidence suggests a broader role in learning. Supporting this view, studies in the hippocampus show that VIP activity increases in novel environments, where its silencing impairs recognition memory and its activation facilitates remapping and spatial learning [88,89]. These findings suggested that VIP cells gate a temporal window for plasticity that underpins learning. We hypothesize that on the novel day, VIP cells initiate learning of the new conditions and image set through stimulus-specific encoding. The emergence of stimulus-specific responses may serve to enhance the salience of unpredicted, and thus potentially behaviorally relevant, sensory information [90,91].

We further propose a functional dissociation between VIP and SST interneurons: VIP cells are recruited during absolute novelty to facilitate learning, shaping activity in both excitatory and SST populations, while SST cells contribute to the storage and maintenance of familiar representations. Notably, the mechanisms triggering the transition in VIP activity on Day 4—whether mediated by local circuits, neuromodulation, or top-down input—warrant further investigation.

VIP cells are concentrated in superficial cortical layers, which receive substantial top-down input from higher-order regions, positioning them to regulate inhibition and disinhibition across local networks [23,92]. It remains to be determined whether VIP Go responses are supported by top-down input across all days or emerge specifically under absolute novelty, with earlier responses arising primarily from local processing. Targeted experiments inhibiting top-down projections to VIP or EXC neurons could clarify this distinction. We predict that disrupting top-down input to VIP cells would impair responses specifically on the novel day and subsequently affect responses on following days, by disrupting learning processes initiated by VIP activation.

### Stable and distinct VIP responses to omissions

VIP interneurons showed robust responses to stimulus omissions. Importantly, omission responsive VIP cells formed a population distinct from Go responsive VIP cells. This observation aligns with a very recent finding showing that, under familiar conditions, VIP neurons encode task-independent information in this paradigm [93]. Our finding therefore raises the possibility that VIP neurons may encode negative prediction errors, such as the absence of an expected stimulus via pathways separate from those used for positive prediction errors (as suggested in [64] with regard to EXC cells), which in turn may contribute to the differential stability of representations at the single-cell level. Whether these populations receive distinct inputs or project to different downstream targets remains an open question for future investigation. Notably, we also observed minimal overlap between omission and Go responsive neurons in both EXC and SST populations, indicating that such pathway segregation may extend beyond VIP cells.

Additionally, VIP responses to omissions were stable across days at the population and at the single-cell level to the same degree of excitatory neurons. This emphasizes the necessity of examining neuronal stability under diverse response conditions; had only image or Go responses been considered, VIP stability might have been overlooked. Whether VIP cells responsive to absolute novelty also exhibit long-term stability remains unknown, as testing this would require a second novel image set on Day 5—an approach not feasible within the current design.

### Excitatory neurons show partial single-cell stability and encode both stimulus-specific and non-specific novelty

We found that excitatory cells exhibited stable population-level responses composed of both stable and unstable responsive cells, with comparable single-cell stability across all three conditions. The findings on image response stability align with prior work showing that only a subset of neurons (∼23%) remains image responsive over five days, while others are newly recruited (∼35%) [47, 50].

With regard to the novelty responses, EXC neurons exhibited both stimulus-specific and non-specific responses to contextual novelty. This finding was unexpected, as prior studies have primarily reported stimulus-specific novelty responses in EXC cells [22,23]. These results suggest the existence of EXC neurons that are selectively responsive to novelty per se. Moreover, the presence of omission responsive EXC neurons, which also formed a distinct population from Go responsive cells, further supports the existence of novelty driven excitatory responses. However, the circuit mechanisms underlying these responses remain unknown —they may arise from local computations within V1, direct top-down modulation of excitatory neurons, or local inhibitory interactions in which a small subset of VIP and SST novelty responsive cells is sufficient to drive the EXC responses.

### Robust population coding despite limited single-cell stability

Our findings prompt further investigation into how population-level decoding remains robust when only a subset of individual neurons exhibit stable responses across days. Recent work demonstrated that a small subset of highly responsive excitatory neurons in V1 can support accurate image decoding, suggesting that sparsely distributed, stable activity may be sufficient for reliable downstream readout [61]. Consistent with this, we observed strong population-level decoding of image changes and omissions, despite only ∼20% of image-responsive excitatory cells remaining stable across days. These findings align with theoretical frameworks proposing that population codes can tolerate variability in individual neurons through mechanisms such as confining drift to the null space of the population code or compensatory plasticity in downstream circuits [34,49,94–99]. Redundant or degenerate representations, where multiple circuit configurations yield similar output, may enable functional robustness in the face of biological variability [100,34].

In this context, the relatively higher stability observed in SST neurons may either contribute directly to population-level robustness or reflect differences in excitatory-inhibitory connectivity [62]. Together, these observations emphasize that representational stability is not solely determined at the single-cell level but likely emerges from population-level organization and circuit architecture. Our results suggest that the relative stability of inhibitory subtypes, particularly SST cells, could add a stabilizing scaffold upon which more dynamic excitatory and VIP responses operate, enabling flexible yet reliable coding across changing conditions. Future computational models and circuit-level experiments should explore how this balance supports robust coding across changing sensory or behavioral conditions.

### Limitations

Several limitations of this study should be acknowledged. First, the task design introduced variability across experimental days. A paradigm with matched stimuli and behavioral context across all days would allow for a more controlled assessment of how specific factors such as stimulus novelty or task engagement modulate response stability. Together with recent studies [101], our findings support the emerging view that response stability, and its underlying mechanisms, must be interpreted considering cell-type identity and specific task conditions.

Second, regarding population response stability, we defined stability using population-averaged response magnitude, the distribution of response magnitudes, and the proportion of responsive neurons across days. While these measures capture key and interpretable aspects of representational consistency, they do not address higher-dimensional population dynamics, which may provide additional insight into how sensory representations are maintained. At the single-cell level, we assessed whether the same neurons remained responsive across days using a binary classification of responsiveness. Future work could incorporate graded or continuous measures to capture more subtle forms of stability.

Third, while our data suggest a stabilizing role for SST neurons, we did not directly test their causal role. Future experiments using targeted perturbations of SST cells across days could determine whether their relative stability supports population-level consistency.

Finally, the changes in activity seen on Day 4 were transient, returning to baseline levels by Day 5. This rapid renormalization is of interest and may be sleep dependent, as sleep has been implicated in memory consolidation for both excitatory and inhibitory neurons [102–104]. However, whether sleep specifically mediates the transition from absolute novelty to familiar responses has not been directly investigated. Incorporating neural recordings during post-task sleep could clarify these dynamics and provide deeper insights into the roles of the different neurons in novelty and familiarity.

## Materials and Methods

### Description of datasets

We analyzed publicly available calcium imaging data from the Allen Brain Observatory Visual Behavior dataset (https://portal.brain-map.org/circuits-behavior/visual-behavior-2p), accessed via the Allen Software Development Kit (AllenSDK). This resource provides Neurodata Without Borders (NWB) files containing preprocessed ΔF/F traces and detailed behavioral metadata. Full experimental details are described in the technical white paper provided by the Allen Institute.

The dataset includes recordings from three transgenic mouse lines targeting specific neuronal populations: excitatory neurons (Slc17a7-IRES2-Cre; Camk2a-tTA; Ai93[TITL-GCaMP6f]), VIP-expressing interneurons (Vip-IRES-Cre; Ai148[TIT2L-GC6f-ICL-tTA2]), and SST-expressing interneurons (Sst-IRES-Cre; Ai148[TIT2L-GC6f-ICL-tTA2]). Imaging was performed using multi-plane two-photon microscopy across four cortical depths (approximately 75, 175, 275, and 375 μm) in two visual cortical areas: VISp and VISl. The provided ΔF/F fluorescence signals and stimulus presentation timestamps were used for all analyses.

To enable longitudinal analyses, we restricted most of our analyses to neurons recorded across all six imaging days. Figure 1 presents results from the full neuronal population to provide an overall view of activity patterns. Figures S1 and S2 show the same analyses from Figure 1, repeated using only the subset of longitudinally tracked neurons, confirming comparable response patterns. All subsequent analyses (Figures 2-5) were conducted exclusively on this longitudinal subset, reducing variability associated with fluctuations in cell sampling across sessions and enabling direct tracking of the same neurons over time.

### Change detection task overview

Mice were trained on a go/no-go visual change detection task. A continuous stream of natural images was presented (250 ms image flashes interleaved with 500 ms gray screens), and mice received a water reward for licking within a brief response window following a change in image identity (Go). The number of repeated image flashes preceding a change was drawn from a geometric distribution, corresponding to a range of 4-11 image presentations (3-9 seconds).

### Experimental schedule

Prior to imaging, mice were trained on the task using a familiar set of eight natural images. During the imaging phase, active behavior sessions were alternated with passive viewing sessions in which no reward spout was present. On days 1-3, mice performed the task with the familiar image set. On day 4, a novel image set was introduced, which remained in use through days 5 and 6. Stimulus omissions were introduced during imaging and occurred on 5% of non-change image presentations across all six days.

### Stimuli evoked neuronal responses

Neuronal calcium traces were aligned to the onset of either image changes or stimulus omissions. For each neuron, responses were averaged across trials, and population averages were computed by averaging across neurons. To quantify image, image change, and omission evoked responses, the mean fluorescence (ΔF/F) was calculated within a 500 ms window following stimulus onset and averaged across trials. Response magnitude was defined as the difference between the response to an image change or omission and the response to the preceding image. To examine how Go responses varied with stimulus repetition, trials were grouped by the number of image repeats preceding the change, and average peak responses were computed for each number of repetitions.

### Fraction of image responsive neurons

To identify image responsive neurons, we analyzed activity during the first four repeated identical image presentations that preceded each image change. A neuron was classified as responsive to a given image if it met three criteria. First, statistical significance: for each trial, the mean activity within a 500ms window following stimulus onset was compared to a null distribution generated by randomly sampling 10,000 time-matched segments from a 5-minute gray screen period preceding the session. A p-value was computed for each image stimulus presentation by the fraction of shuffled responses that exceeded the mean stimulus response. A neuron was considered significant if at least 25% of trials for that image yielded p < 0.05.

The second criteria was magnitude threshold: the neuron’s mean response had to exceed the 90% confidence interval of the shuffled gray screen distribution, ensuring a response magnitude above spontaneous baseline variability. The third criteria was frequency consistency: to assess stimulus locked periodicity in the responses, a fast Fourier transform was applied to each trial’s calcium trace. The amplitude at 1.33 Hz-corresponding to four image repetitions within a 3-second window was extracted. A null distribution was generated by applying the same analysis to overlapping 3-second segments of gray screen data. A neuron was considered responsive in the frequency domain if at least 10% of trials showed a significant peak at 1.33 Hz (p < 0.05), reducing false positives. The fraction of responsive neurons was computed as the number of neurons meeting all three criteria divided by the total number of recorded neurons across all six experimental days.

### Fraction of Go responsive neurons

To identify neurons responsive to image changes (Go), activity was analyzed during each trial and focused on the two repeated identical image presentations followed by a changed image. Neurons were classified as Go responsive if they showed a consistent increase in activity following the image change. For each neuron, the average activity was computed at three time points: (1) the image preceding the image before the Go, (2) the image immediately preceding the Go (identical image), and (3) the Go image (the changed stimulus). To determine whether the Go evoked a significant increase in activity, we first quantified baseline variability by calculating the difference between the responses to the two identical images preceding the Go. The 95th percentile of this distribution was used as a threshold for detecting meaningful changes in activity. A trial was considered Go responsive if the difference between the Go response and the immediately preceding image exceeded this threshold. A neuron was classified as Go responsive if at least in 10% of trials the response difference exceeded the threshold, ensuring consistent activation beyond spontaneous fluctuations.

### Fraction of omission responsive neurons

Neurons were classified as omission responsive based on increased activity during the absence of an expected image stimulus. For each neuron, responses were analyzed across all omission trials by comparing activity during the omission window to the preceding image presentation. Classification was based on three criteria. First, to ensure responses exceeded spontaneous fluctuations, activity during omissions was compared to a shuffled distribution generated from 10,000 resampled gray screen traces. A neuron was considered omission responsive if at least 10% of trials showed omission responses greater than expected by chance (p < 0.05). Second, increase in activity during the omission and third, the difference between omission and pre-omission responses had to be greater than zero.

### Quantification of response stability at the population and single-cell levels

To assess population-level stability, response magnitudes were calculated for each stimulus condition (image, Go, or omission) by averaging responses across trials for neurons classified as responsive to that condition on a given day. We compared the fraction of responsive cells, as well as the mean and distribution of response magnitudes, across days.

For single-cell-level stability, we quantified the proportion of neurons that remained responsive to the same stimulus across multiple days. Specifically, for each stimulus, we computed the number of neurons that were responsive on all three consecutive days (analyzed for the familiar and novel image sets) and divided it by the total number of neurons responsive to that stimulus on at least one of those days. A similar analysis was conducted over two-day intervals. For image responses, stability was computed separately per image and then averaged across all images in the set. All percentages were computed separately for each mouse.

To determine whether the observed percent of stable cells exceeded chance levels, we generated a null distribution by randomly permuting the identity of responsive neurons across days (10,000 iterations). For each shuffle, we computed the percentage of neurons responsive on all three days, resulting in a null distribution of response overlap. Observed stability was considered significant if it exceeded the 99th percentile of the null distribution.

### Decoding image change occurrence

To determine whether single-trial population activity encodes image changes, we trained a logistic regression classifier to distinguish neuronal responses across successive image presentations. Each trial was divided into three periods: image 1 (the image preceding the image before the Go), image 2 (the image before the Go), and image 3 (the Go image). For each period, the average activity within a 500 ms window following stimulus onset was computed for all neurons. The classifier was trained on each day to differentiate between two conditions: image 1-image 2 (class 0, no change) and image 2-image 3 (class 1, image change). Data were randomly shuffled and split into training and test sets. Model performance was evaluated using 8-fold cross-validation, and classification accuracy was defined as the mean accuracy across folds. To establish chance-level performance, we generated a null distribution by permuting test set labels and computing classifier accuracy on the shuffled data, yielding an expected baseline accuracy near 50%. All model training and evaluation were performed independently for each of the six experimental days.

### Decoding omission occurrence

To determine whether single-trial population activity encodes the occurrence of stimulus omissions, we employed a classification approach similar to that used for decoding image changes. Each omission trial included two consecutive image presentations followed by an omitted stimulus. Average neuronal activity was computed in 500ms windows following each image onset and during the omission period.

### Stimulus specific and non-specific novelty responses

To examine whether responses to contextual novelty are stimulus-specific (convey the identity of the stimulus) or non-specific, neurons were categorized into three groups based on their responsiveness to images and to image changes. The first group, termed “*Image + Go Responsive*” included neurons that were both responsive to specific images (as defined in the section “Fraction of image responsive neurons”) and Go responsive (as defined in the section “Fraction of Go responsive neurons”). The second group, termed “*Go-Only Responsive*” included neurons responsive to the Go but did not show responses to individual images. The third group, “*Go-Nonresponsive*” included neurons that were not classified as Go responsive. For each neuron, the average response to the image immediately preceding the Go image (Before Go image) and to the Go image was computed and the response difference was calculated within each group. This allowed us to evaluate the relative contribution of each group to the overall Go response.

To further assess stimulus specificity in “*Image + Go Responsive*” neurons (group 1), neuronal responses were analyzed across four distinct conditions: (1) Before Go Preferred: Response to a pre-Go image when the image was one to which the neuron was previously classified as responsive (termed neuron’s preferred image). (2) Before Go Non-Preferred: Response to a pre-Go image that was not among the neuron’s preferred images (non-preferred image). (3) Go Preferred: Response to the Go image when the image was a preferred image. (4) Go Non-Preferred: Response to the Go image when the image was non-preferred. Preferred and non-preferred images for each neuron were defined based on image responsiveness (during repeated image presentation before image changes), as outlined in the section “Fraction of image responsive neurons”. We compared response magnitudes across these four conditions to determine whether neurons in group 1 exhibited stimulus-specific responses during image changes.

### Quantification of neuronal firing rates from electrophysiological data

To complement our two-photon calcium imaging dataset, we analyzed extracellular spiking activity from the Allen Institute’s publicly available Visual Behavior Neuropixels dataset (https://portal.brain-map.org/explore/circuits/visual-behavior-neuropixels). Firing rate traces were extracted for neurons recorded in visual areas VISp and VISl to align with the regions analyzed in the calcium imaging analysis. Neurons were categorized into four putative cell classes: EXC, SST, VIP and PV populations. SST and VIP neurons were identified through optotagging, following the procedure described in the Allen SDK documentation (https://allensdk.readthedocs.io/en/latest/_static/examples/nb/ecephys_optotagging.html). Neurons not tagged optogenetically were classified based on waveforms: units with waveforms duration shorter than 0.4 ms were considered putative PV neurons (fast-spiking inhibitory), while those with longer waveforms durations were classified as putative excitatory neurons. Only units that met stability criteria were used. The dataset consisted of recordings from both male and female mice, specifically Sst-IRES-Cre/wt;Ai32(RCL-ChR2(H134R)_EYFP)/wt mice and Vip-IRES-Cre/wt;Ai32(RCL-ChR2(H134R)_EYFP)/wt mice.

Mice underwent two acute recording days, with no attempt to match individual neurons across days. On the first recording day, visual stimuli were drawn from a familiar set of eight natural images that the mice had been previously trained on. On the second day, a novel image set was introduced. This set consisted of six previously unseen (“non-shared”) images and two familiar (“shared”) images that had also been shown during the first day. Each recording day included an active session followed by a passive viewing session. The behavioral task was identical to that used in the calcium imaging dataset.

### Stimuli evoked neuronal responses

Peri-stimulus time histograms (PSTHs) were computed by binning spiking activity into 1 ms bins aligned to stimulus onset. Mean firing rates were obtained by dividing spike counts by the bin size and number of trials. Responses were calculated separately for each cell type under the following conditions: Day 1 (familiar) active and passive sessions for shared and non-shared images, and Day 2 (novel) active and passive sessions for shared and non-shared images. For visualization purposes, PSTHs were smoothed by averaging across every 5 consecutive bins in the time-course plots.

Latency to peak firing was calculated for each neuron within a 150 ms window following stimulus onset, separately for the image preceding the Go (BG) and the Go image, across all Go trials. Only neurons with an average of at least one spike per stimulus presentation across trials were included in the analysis.

### Statistics

All statistical analyses were conducted using custom Python scripts and GraphPad Prism, with significance defined at α = 0.05. Analyses were performed at either the neuron or mouse level. One-way or two-way ANOVAs were used to assess effects of between-subject factors such as cell type, and within-subjects factors such as time, condition (stimulus type) or group as appropriate. Comparisons between two groups were assessed using paired or unpaired two-tailed t-tests, depending on whether measurements were repeated within subjects. When data included repeated measurements from the same mice across days or conditions, we used repeated-measures ANOVAs. Post hoc comparisons were performed using Tukey’s, Dunnett’s, Sidak’s, or Fisher’s LSD tests selected based on the structure of the data and the experimental hypothesis. Factors such as image preference (preferred vs. non-preferred), behavioral timing (before vs. during Go), and stimulus familiarity (familiar vs. novel) were included in statistical models when relevant. To assess changes in the distribution of response magnitudes across days, we used pairwise Kolmogorov-Smirnov tests comparing Day 1 to subsequent days, with Bonferroni correction for multiple comparisons. All test statistics, degrees of freedom, and p-values are reported in the figure legends. Significance levels were reported as ns p>0.05, *p < 0.05, **p < 0.01, ***p< 0.001, ****p<0.0001. All statistical tests were two-sided.

## Resource availability

### Lead contact

Further information and requests should be directed to Karni Lev Bar-Or.

### Data and code availability

This paper analyzes existing, publicly available data which is available via the AllenSDK at: https://allensdk.readthedocs.io/en/latest/visual_behavior_optical_physiology.html and: https://allensdk.readthedocs.io/en/latest/visual_behavior_neuropixels.html. Original code will be deposited and publicly available as of the date of publication. Any additional information required to reanalyze the data reported in this paper is available from the lead contact upon request.

## Acknowledgments

We thank Ruthy Lev Bar-Or and Inna Slutsky for thoughtful and insightful comments on the study questions and manuscript. We are also grateful to the members of the Slutsky lab, as well as Daniel Ekeltchik, Ran Darshan, and Lee Susman, for their valuable feedback on the manuscript. We thank Neuromatch Academy and Sajjad Rezvani for facilitating the collaboration among the authors. K.L.B.O. acknowledges support from the Sieratzky Institute for Advances in Neuroscience through a fellowship of excellence for outstanding Ph.D. students.

## Author contributions

Conceptualization, K.L.B.O; Software & formal analysis, K.L.B.O, V.S.K; Writing, K.L.B.O; Methodology: K.L.B.O, V.S.K, D.G.W

## Supporting information

**S1 Fig.**
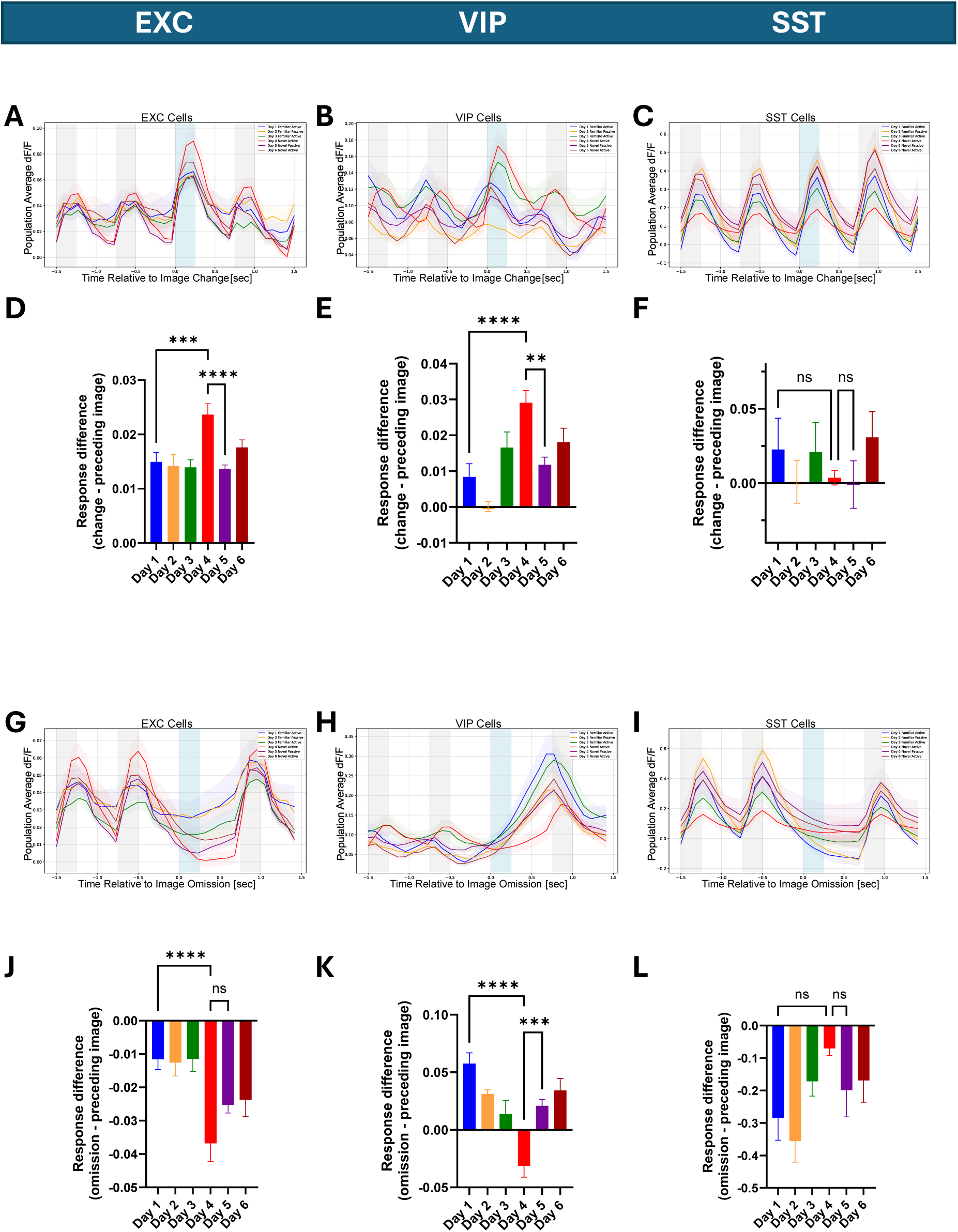
Cell type-specific responses to different novelty conditions in common cells tracked across all six days. Related to Fig 1. (A-C) Population averaged responses to image presentations (gray) and image changes (Go, blue) for cells tracked across all six experimental days. Shown for EXC (n=694 cells), VIP (n=177 cells), and SST (n=53 cells). (D-F) Go response magnitude (difference between image change and preceding image). EXC and VIP cells show increased responses on Day 4 (one-way ANOVA followed by Dunnett’s post hoc comparing all days to Day 4; EXC: F(5, 4158) = 5.6, p < 0.0001, all multiple comparisons were significant, p < 0.001; VIP: F(5, 1056) = 8.8, p < 0.0001, multiple comparisons, p ≤ 0.001; SST: F(5, 312) = 0.7, p = 0.6, multiple comparisons were ns). (G-I) Population average responses to stimulus omissions (blue) across days for cells tracked across all six experimental days. (J-L) Omission response magnitude (difference between omission and preceding image). VIP responses are reduced on Day 4 (one-way ANOVA followed by Dunnett’s post hoc comparing all days to Day 4; EXC: F(5, 4158) = 6.1, p < 0.0001, multiple comparisons showed a significant difference for Day 1, p<0.0001; VIP: F(5, 1056) = 10.9, p < 0.0001, all multiple comparisons were significant, p < 0.001; SST: F(5, 312) = 2.6, p = 0.02, multiple comparisons were ns except for the comparison with Day 2). Error bars: ± SEM across neurons. ns p>0.05, *p < 0.05, **p < 0.01, ***p< 0.001, ****p<0.0001.

**S2 Fig.**
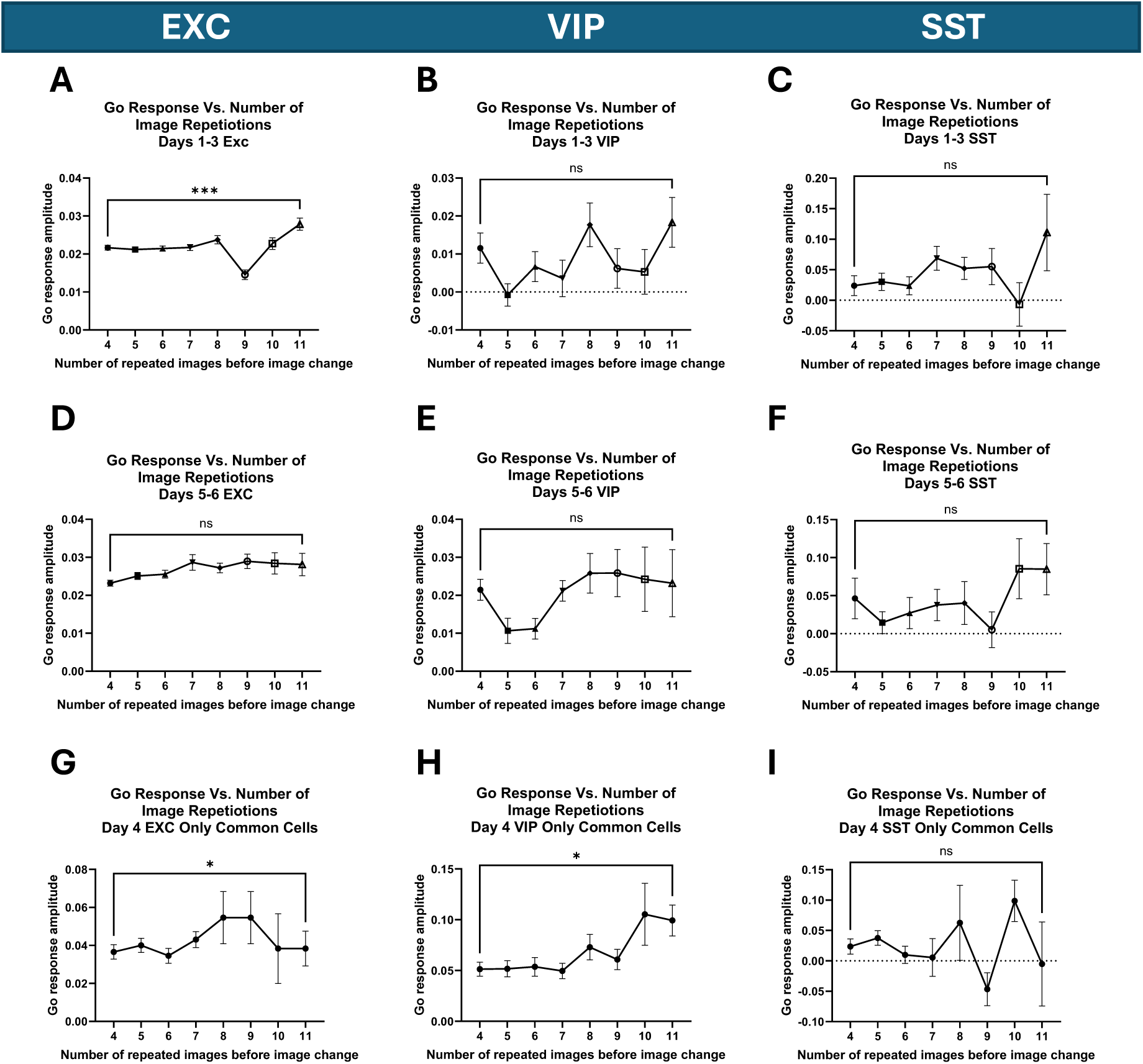
Dependence of Go response on number of image repetitions. Related to Fig 1. (A-C) Go response as a function of the number of image repetitions preceding the image change on Days 1-3. EXC neurons showed increased responses when the image change was preceded by more repetitions, indicating greater unexpected event. No significant modulation was observed in VIP or SST cells (unpaired t-test; EXC: t(5998) = 3.6, p = 0.0003; VIP: t(798) = 0.9, p = 0.4; SST: t(198) = 1.3, p = 0.2) (D-F) Same analysis for Days 5-6 shows no significant modulation (EXC: t(5998) = 1.6, p = 0.1; VIP: t(798) = 0.2, p = 0.8; SST: t(198) = 0.9, p = 0.4). (G-I) Same analysis for Day 4, restricted to cells tracked across all six days. A similar trend to the one seen in Figure1 is observed, Go responses are larger following more repetitions in EXC and VIP cells (Wilcoxon matched-pairs signed-rank test; EXC: W = 25985, p = 0.01; W = 1699, p = 0.04; SST: W = -97, p = 0.7). Error bars: ± SEM across neurons. ns p>0.05, *p < 0.05, **p < 0.01, ***p< 0.001, ****p<0.0001.

**S3 Fig.**
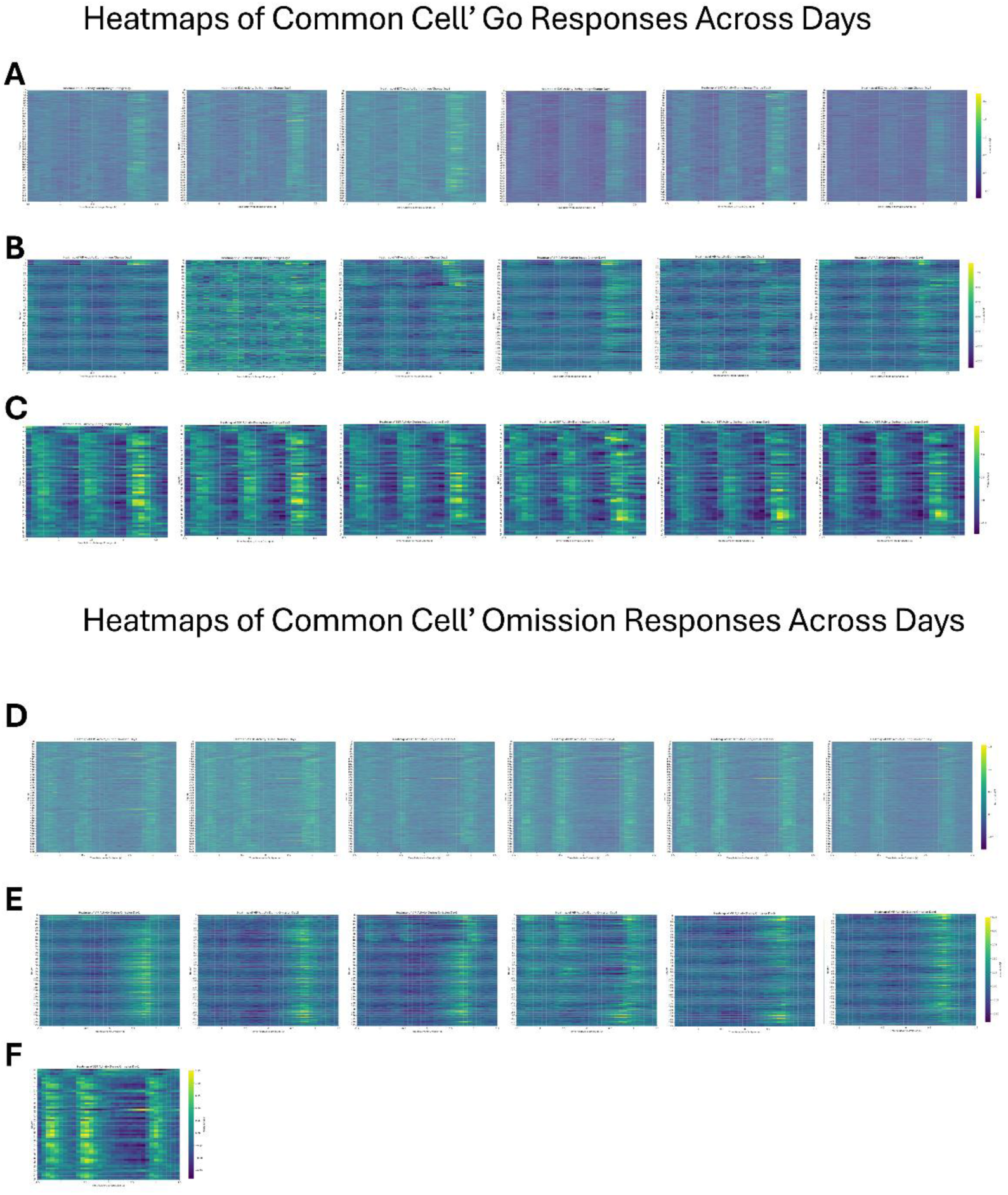
Longitudinal tracking of Go and omission responses in individual neurons. Related to Figs 3-4. (A-C) Normalized ΔF/F (per day for visualization) responses of the same cells across days during two repeated image presentations followed by an image change (Go). Responses shown for (A) EXC, (B) VIP, and (C) SST neurons. (D-F) Normalized ΔF/F responses of the same cells across days during two image presentations followed by an omission, and then another image. Responses are shown for (D) EXC, (E) VIP, and (F) SST neurons.

**S4 Fig.**
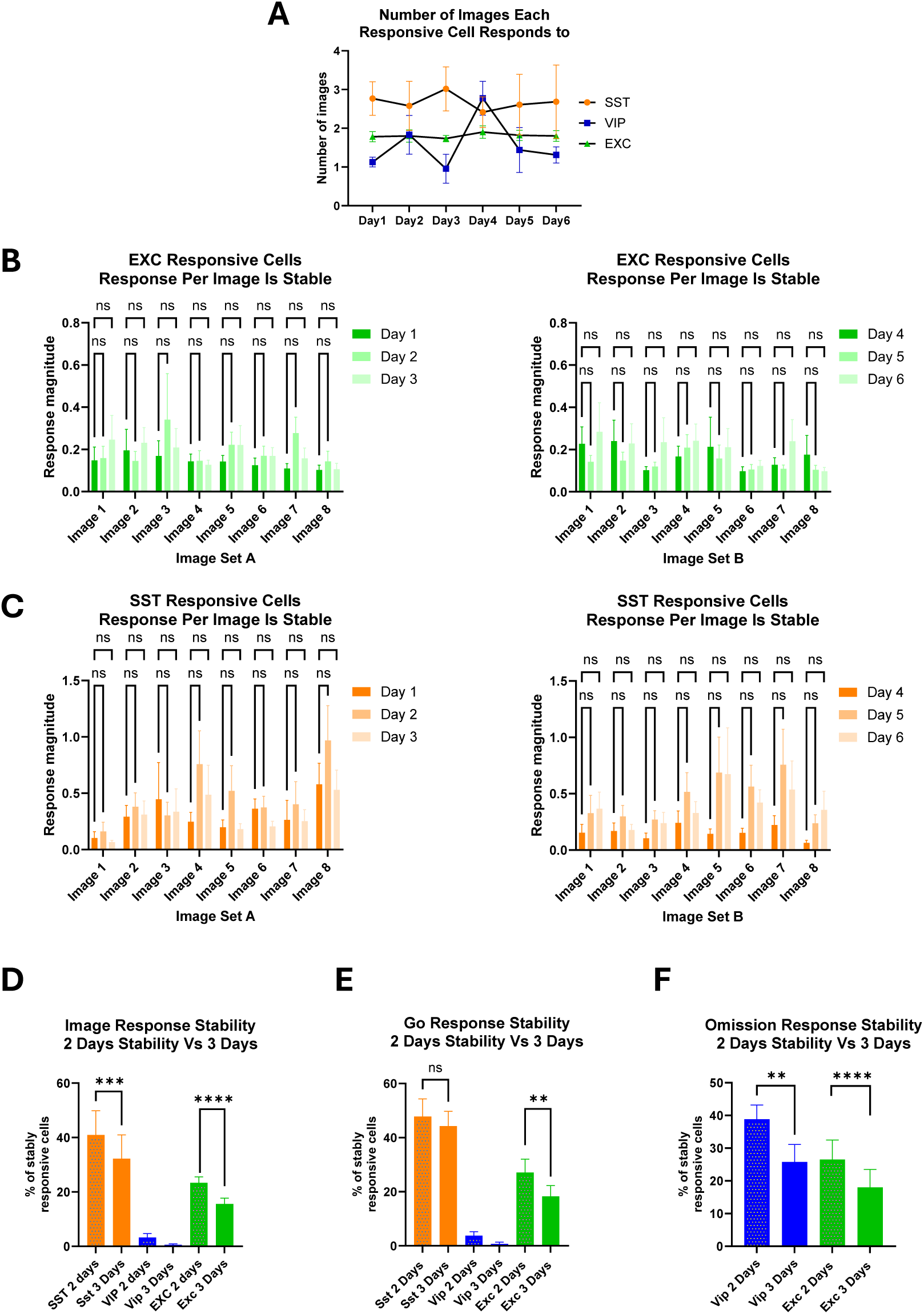
Cell type-specific stability of neuronal responses across days. Related to Figs 2-4. (A) Average number of images to which image responsive neurons responded per day, related to Figure2 (two-way RM ANOVA followed by Tukey’s multiple comparisons; day F(2, 18) = 1, p = 0.3, all comparisons between days were ns). (B-C) Response magnitudes of image responsive neurons remained stable across days for each image. (B) EXC cells: Left: Days 1-3. For each image, the response magnitude of the neurons classified as image responsive on that day remained consistent across days (two-way ANOVA; day: F(2, 1469) = 1, p = 0.3. Dunnett’s post hoc comparisons showed no significant differences between days for any image, all p > 0.1). Right: Days 4-6. Similarly, responses were stable across days (two-way ANOVA; day: F(2, 1848) = 2, p = 0.2, post hoc, p > 0.4). (C) SST cells: Left: Days 1-3. Responses were stable across days (two-way ANOVA; day: F(2, 443) = 0.4, p = 0.9, post hoc, p > 0.05). Right: Days 4-6. Responses remained stable across days (two-way ANOVA; day: F(2, 427) = 4, p = 0.02, post hoc, p > 0.1). (D-F) The proportion of stable neurons across two consecutive days was higher than across three-day intervals, as expected (paired t-test; image: EXC: t(5)=32, p<0.0001; SST: t(2)=33, p<0.001; Go: EXC: t(5)=6, p<0.01; SST: t(2)=1, p=0.3; omission: EXC: t(5)=12, p<0.0001; VIP: t(3)=7, p<0.01). Error bars: (A, D-F) ± SEM across mice; (B-C) ± SEM across neurons. ns p>0.05, *p < 0.05, **p < 0.01, ***p< 0.001, ****p<0.0001.

**S5 Fig.**
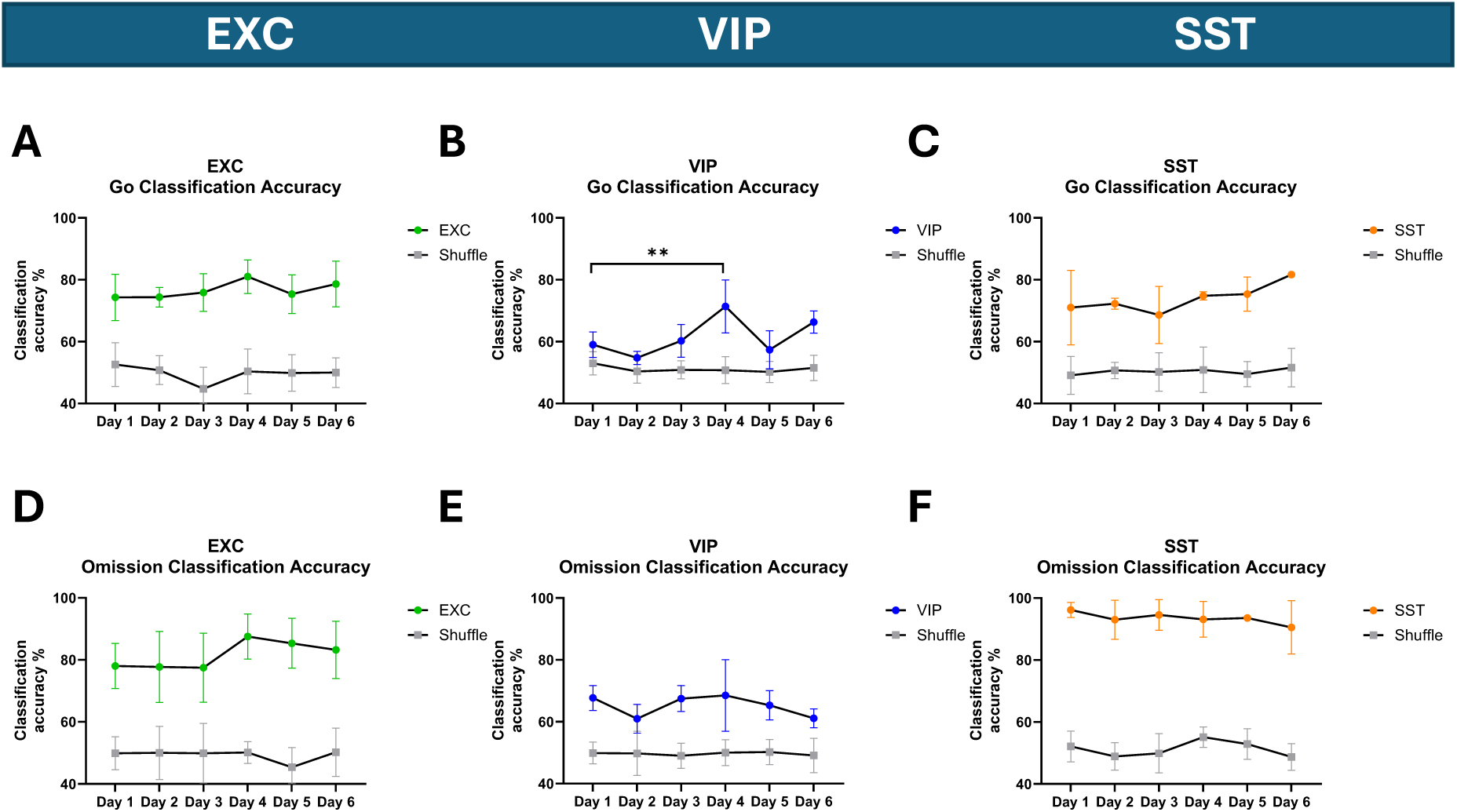
Population decoding of image changes and omissions is stable across days. Related to Figs 2-4. (A-C) Logistic regression decoding of image change versus no change, from the population code, showed stable accuracy across days and exceeds chance level for all cell types. Models were trained and tested separately for each day using activity from neurons recorded on all six days during two repeated image presentations (no change) and during the image preceding and including a change (Go) (two-way RM ANOVA followed by Tukey’s multiple comparisons; EXC: effect of model, F(1, 84) = 426, p < 0.0001, effect of day, F(5, 84) = 1, p = 0.3; VIP: model, F(1, 72) = 104, p < 0.0001, day, F(5, 72) = 6, p < 0.0001. Post hoc showed significant difference between Day1 and Day4; SST: model, F(1, 60) = 168, p < 0.0001, day, F(5, 60) = 1, p = 0.3. For all cell types, decoding accuracy exceeded chance (shuffled data) on all days). (D-F) Logistic regression decoding of omission versus no omission stimuli, from the population code, also remained stable and above chance across days. Models were trained and tested separately for each day using activity from neurons recorded on all six days during two repeated image presentations (no omission) and during the image preceding and including an omission. (two-way RM ANOVA followed by Tukey’s multiple comparisons; EXC: model, F(1, 84) = 371, p < 0.0001; day, F(5, 84) = 1, p = 0.4; VIP: model, F(1, 72) = 151, p < 0.0001; day, F(5, 72) = 1, p=0.3; SST: model, F(1, 60) = 753, p < 0.0001; day, F(5, 60) = 1, p = 0.5. For all cell types, decoding accuracy exceeded chance (shuffled data) on all days). Error bars: ± SEM across mice. ns p>0.05, *p < 0.05, **p < 0.01, ***p< 0.001, ****p<0.0001.

**S6 Fig.**
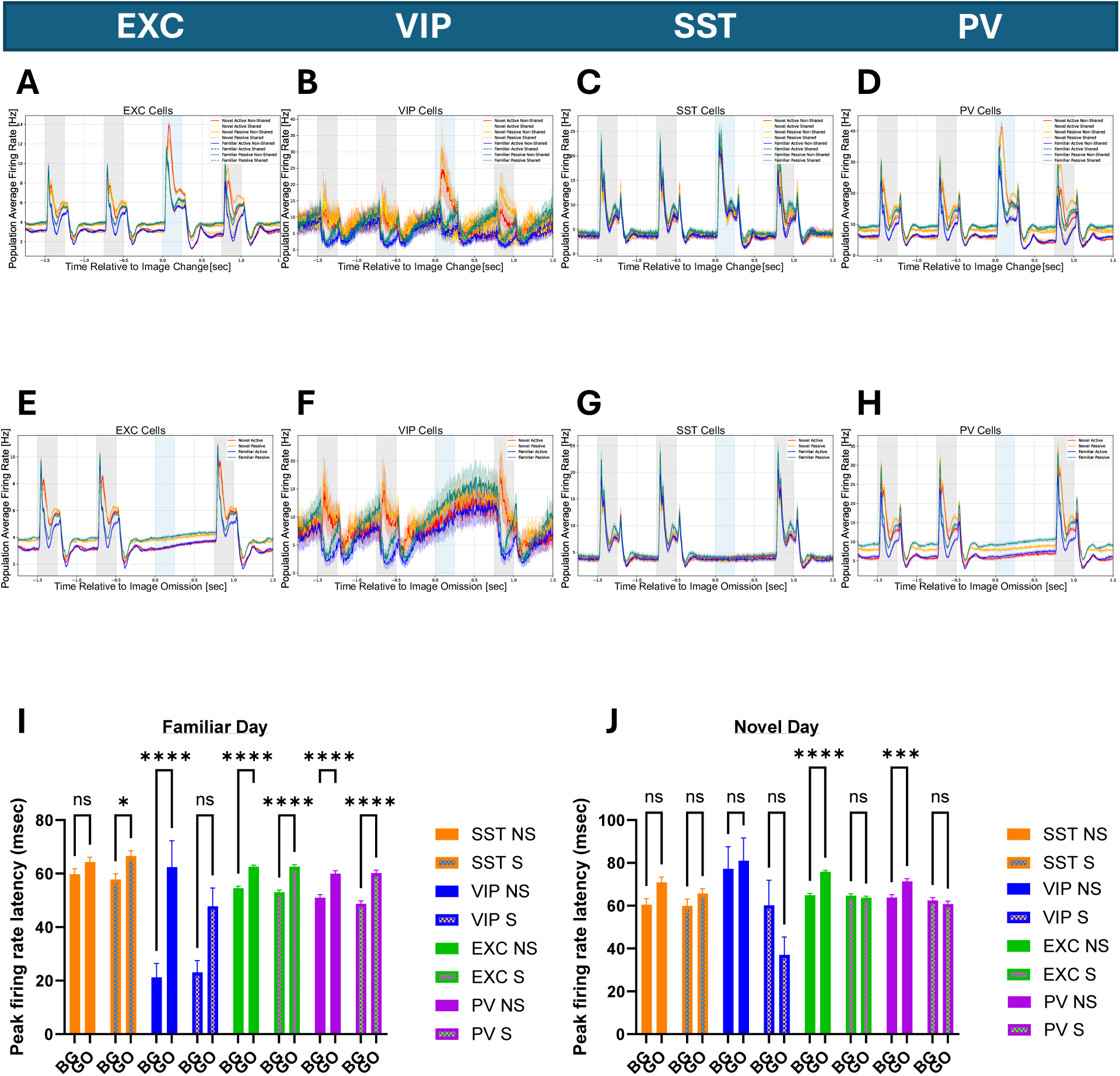
Firing rate dynamics in familiar and novel conditions. Related to Fig 6. (A-D) Population average responses to images (gray) and image changes (Go, blue) on both the familiar and novel days, shown separately for active and passive sessions (EXC: 3842 ± 317 neurons from n=27 mice, SST: 220 ± 26 neurons from n=19 mice, VIP: 17 ± 5 neurons from n=7 mice, PV: 759 ± 105 neurons from n=27 mice). (E-H) Population average responses to image omissions (blue) across all sessions. VIP neurons show increased activity during omissions. (I) On the familiar day, EXC, VIP and PV cells exhibited increased response latency during Go stimuli compared to the preceding image for non-shared (NS) images (two-way ANOVA; main effect of Go, F(1, 7458) = 73, p<0.0001, Sidak’s post hoc: BG vs. GO NS, p<0.0001 for EXC and PV and VIP). (J) On the novel day, PV and EXC cells showed increased response latency during Go stimuli compared to the preceding image only for non-shared images. VIP neurons showed a trend toward reduced latency during Go for shared images (two-way ANOVA; main effect of Go, F(1, 7239) = 0.5, p=0.5, Sidak’s post hoc: BG vs. GO NS, p<0.001 for EXC and PV). Error bars: ± SEM across neurons. ns p>0.05, *p < 0.05, **p < 0.01, ***p< 0.001, ****p<0.0001.

